# ALS-associated FUS mutation reshapes the RNA and protein composition of Stress Granules

**DOI:** 10.1101/2023.09.11.557245

**Authors:** Davide Mariani, Adriano Setti, Francesco Castagnetti, Erika Vitiello, Lorenzo Stufera Mecarelli, Gaia di Timoteo, Andrea Giuliani, Eleonora Perego, Sabrina Zappone, Nara Liessi, Andrea Armirotti, Giuseppe Vicidomini, Irene Bozzoni

## Abstract

Stress Granules (SG) formation is a cellular protection mechanism, constituting a storage for untranslated mRNAs and RNA-binding proteins (RBPs); however, these condensates can turn into pathological aggregates, related to the onset of neurodegenerative diseases like Amyotrophic Lateral Sclerosis (ALS). This transition towards cytotoxic inclusions is triggered by ALS-causative mutations in the RBP FUS, which lead to its cytoplasmic mis-localization and accumulation in SG. Here, we describe the SG transcriptome in a neural context and describe several features for RNA recruitment in SG. We demonstrate that SG dynamics and RNA content are strongly modified by the incorporation of mutant FUS, switching to a more unstructured, AU-rich SG transcriptome. Moreover, we show that mutant FUS, together with its protein interactors and their target RNAs, are responsible for the reshaping of the mutant SG transcriptome with alterations that can be linked to neurodegeneration. Therefore, our data give a comprehensive view of the molecular differences between physiological and pathological SG in ALS conditions, showing how FUS mutations impact the RNA and protein population of these condensates.

## INTRODUCTION

A well-defined intracellular compartmentalization is vital for eukaryotic cells, with membrane-bound organelles physically separating specialized molecular processes from the rest of the cellular environment. Alongside these structures, a large number of membrane-less organelles (MLOs) has been identified in recent years as crucial in the regulation of gene expression and metabolic activity of cells^1–3^. Lacking a lipid bilayer, these assemblies are separated from the neighboring cytoplasm or nucleoplasm by liquid-liquid demixing, conferring them stability, but also dynamic properties of exchange with the molecular surroundings^4–6^. The liquid-like behavior of MLOs, mainly constituted by diverse RNAs and RNA binding proteins (RBPs), results from the summation of several weak interactions involving its constituents^7^.

RNA is a pivotal component of such condensates, since the intrinsic characteristics of individual transcripts, such as length, structure and nucleotide composition, are sufficient to influence the number of RNA-RNA and RNA-protein interactions within a granule, thereby affecting its physical properties^8–11^. Moreover, it has been observed that the presence of Intrinsically Disordered Regions (IDRs) in RBPs is a molecular determinant of their tendency to phase-separate, both in vitro and in living cells^12^; IDRs are often coupled to canonical RNA binding domains within the same protein, providing the foundation for establishing multivalent interactions with RNAs and other proteins, thus leading to self-assembling, separated granules^13,14^.

The formation of cytoplasmic Stress Granules (SG) in response to variations in cell homeostasis stimuli is a primordial protection mechanism^15^. SG result from the condensation of untranslated, stalled mRNAs and several IDR-containing RBPs, functioning as a temporary reservoir and being readily disassembled as protein translation is reactivated after stress^16^. SG assembly is a multi-step event, where initial phase separation of individual ribonucleoparticles leads to the coalescence of stable *cores*, surrounded by a liquid, quickly remodeled *shell*^17^.

Post-mitotic cells like neurons are highly susceptible to dysregulation of SG assembly and disassembly^18–20^. Mutations in numerous protein components of these condensates are causative of the formation of permanent, insoluble protein-RNA assemblies; these aggregates have been causatively linked to the pathogenesis of neurodegenerative diseases and constitute a hallmark of several motor-neuron diseases, such as Amyotrophic Lateral Sclerosis (ALS)^21,22^.

A paradigmatic example is represented by FUS/TLS, whose mutations account for 4% of familial ALS cases^23^. The protein embeds a IDR which is responsible for self-assembly and spontaneous aggregation^24–26;^ ALS-linked mutations are clustered in the Nuclear Localization Sequence (NLS) reducing its nuclear reimport and resulting in an aberrant cytoplasmic accumulation and subsequent recruitment in SG^27,28^. As mutant FUS accumulates in the cytoplasm of motor-neurons, it triggers the activation of major stress pathways and impacts SG number and dynamics, finally leading to an increased vulnerability of these cells to harmful stimuli^29^.

In this work, we focused on the RNA content of SG in a neuronal-like system, demonstrating that neuronal RNAs are specifically enriched in these condensates in response to oxidative stress. We show that the expression of the mis-localized FUS^P525L^ mutant is sufficient to impact the dynamics of SG and to strongly modify their transcriptome by enriching AU-rich RNAs, at the expenses of GC-rich transcripts. The modification of the nucleotide composition caused by the presence of mutant FUS reduces the global structuration propensity of SG RNAs, by recruiting poorly structured transcripts. Moreover, we show that the reorganization of the SG transcriptome in FUS^P525L^ conditions affects specific classes of RNAs linked to neurodegenerative processes. Finally, we show that the presence of FUS^P525L^ also impacts on the SG proteome; thus, while the direct binding of mutant FUS to RNA is not sufficient to explain the altered engagement of transcripts into SG, the differential recruitment of FUS-interacting RBPs contributes to the aberrant RNA composition of pathological granules.

## RESULTS

### Characterization of the RNA composition of neural-like SG

To investigate the RNA composition of neuronal Stress Granules (SG), we adapted the purification protocol described by Khong et al.^30^ (Extended Data Fig. 1A) on a SK-N-BE neuroblastoma cell line expressing the SG core protein G3BP1 fused to a GFP tag (SK-GFP-G3BP1). Fluorescence microscopy showed that upon oxidative stress these cells display GFP-G3BP1 cytoplasmic foci, confirming that SG were correctly assembled (Extended Data Fig. 1B). Western Blot analysis confirmed the specific enrichment of the endogenous SG marker TIAR in the IP fraction of SK-GFP-G3BP1 only in stress conditions compared to unstressed control, suggesting that the SG are correctly purified (Extended Data Fig. 1C). Moreover, after RNA extraction, qRT-PCR was performed on targets previously identified as enriched or depleted in granules of U2OS cells^31^. AHNAK, CENPF, HUWE1 and TRIO were selected as positive controls and displayed a strong enrichment in the IP fraction under stress induction, while the negative controls ATP5O, COX8A and LGALS1 were not precipitated (Extended Data Fig. 1D).

After these controls, we generated four replicates of purified SG and corresponding total RNA inputs and performed total RNA sequencing. Multidimensional scaling (MDS) plot of leading fold change (Extended Data Fig. 1E) and Pearson’s correlation matrix (Extended Data Fig. 1F) confirmed the expected clusterization of the samples. We proceeded to RNA-Sequencing analysis comparing SG-enriched (SG-enr) versus Input samples (INP). Among the 12318 RNAs detected in the system (>1 FPKM in INP or SG-enr) we identified 1816 enriched (log_2_FC > 1 and FDR < 0.05) and 2939 depleted (log_2_FC < -1 and FDR < 0.05) transcripts in the SG core (Fig. 1A, Supplementary Table 1). As expected, most of the RNAs that we identified as enriched in SG were mRNAs (85% of the enriched fraction) and among the non-coding fraction (15%) the most represented category was long non-coding RNAs (9%) (Extended Data Fig. 1G).

**Figure 1.**
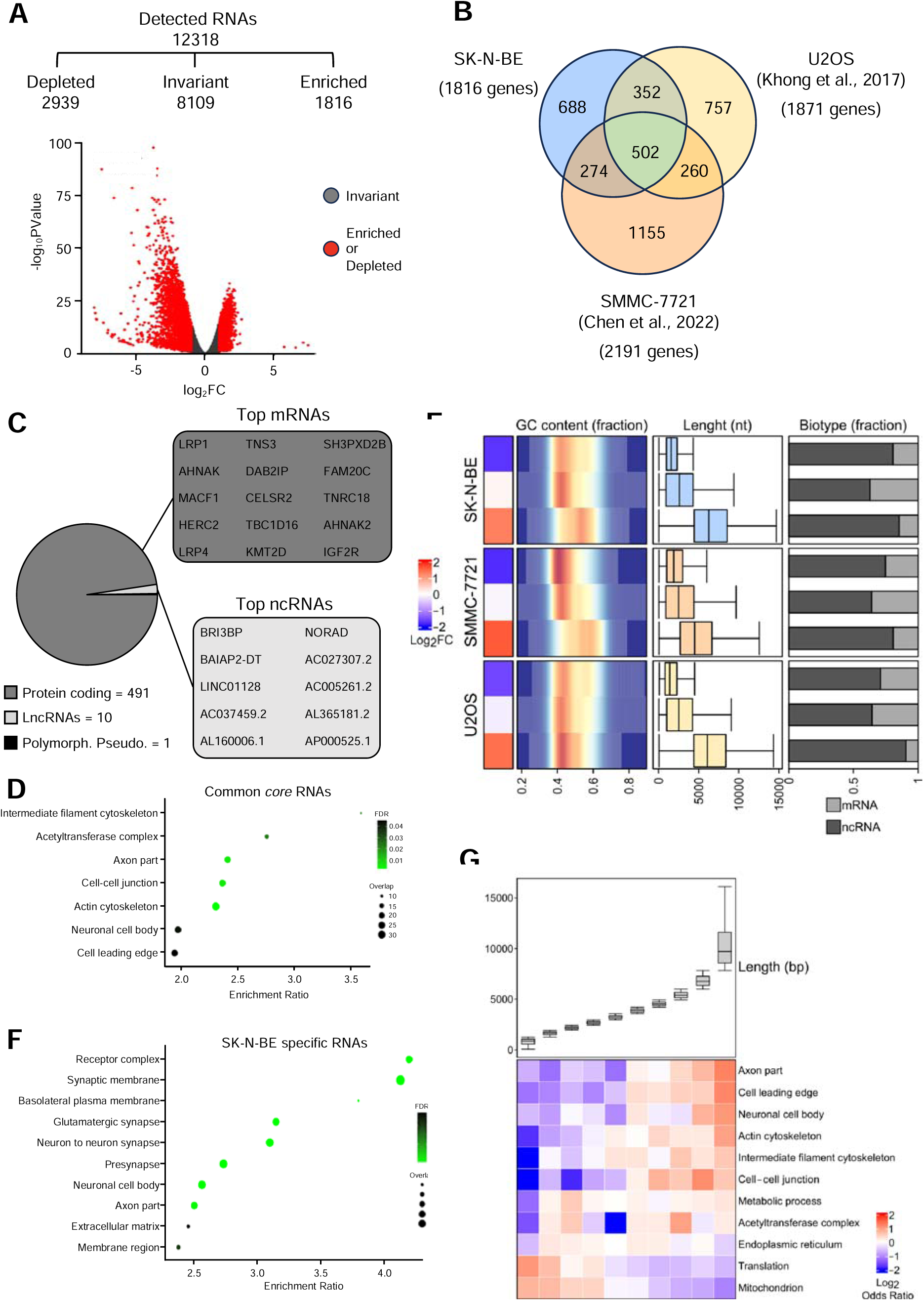
**A) Upper panel.** Summary of the amount of RNAs detected in RNAseq of Stress Granule (SGs) purification experiment, and their partition among enriched, invariant and depleted groups. **Lower panel**. Volcano plot showing the log2 fold change (SGenr/Input) and the -log10 p-value of SG enrichment for each RNA detected. Significantly enriched and depleted RNAs (FDR < 0.05, |logFC| > 1) or invariant RNAs are indicated by red or gray dots, respectively. **B)** Venn diagram depicting the overlap between SG-enriched RNAs in neuroblastoma (SK-N-BE, this study), osteoblastoma (U2OS) and endocervical adenocarcinoma (SMMC-7721) cell lines. **C) Left panel.** Pie chart depicting the fraction of protein coding and non coding RNAs in the SG common core. **Right panel**. Set of the 15 top enriched mRNAs and all the ncRNAs enriched the SG common *core*. **D)** Dotplot depicting GO over-represented categories in the SG common core. Only significant categories (FDR < 0.05) of Cellular Component database were depicted. X-axis represents category enrichment score while y-axis reports the GO category description. Dot size represents the amount of RNAs in the analyzed group that overlap the category, while color code (green) reports the significativity of the enrichment. **E)** Heatmap depicting the association of RNA features with SG enrichment in SK-N-BE (upper panel), SMMC-7721 (middle panel) and U2OS (lower panel) cell lines. For the defined transcripts group («depleted», «invariant» and «enriched») of each cell line the following characteristics were described: median log2FC (heatmap); GC content (density plot); RNA length (nt,boxplot); transcripts biotypes fraction (barplot). **F)** Dotplot depicting GO over-represented categories of the SK-N-BE SG specific RNAs. Graph features are the same as panel D. **G)** Heatmap (lower panel) depicting the relationship between RNA length and GO categories enriched in “SG common core”. The 10 transcripts groups represented were defined by length-based stratification of the transcriptome. The log2 Odds Ratio depicted in the heatmap shows the depletion (blue) or the enrichment(red) of each GO category in the defined group compared to the remaining fraction of the transcriptome. Boxplot (upper panel) represents the distributions of RNAs length for the 10 transcripts groups.

In order to identify the discriminating features of the obtained neuroblastoma cells SG transcriptome, we compared it to the ones of osteoblastoma (U2OS)^31^ and endocervical adenocarcinoma (SMMC-7721 cell line)^32^ (Supplementary Table 1). We observed a strong correlation of the enrichment scores (log_2_FC) among all the three datasets (Extended Data Fig. 1H), indicating that the methodology is highly reproducible even when applied in different systems. When comparing the SG transcriptomes, we identified a common *core* consisting of 502 RNAs (Fig. 1B). These transcripts are mostly mRNAs (Fig. 1C) and are enriched in Gene Ontology (GO) categories related to cytoskeletal and structural components, including neuronal-specific factors (Fig. 1D, Extended Data Fig. 1I, Supplementary Table 2).

Noteworthy, neuronal transcripts, even if less represented in the transcriptomes of the other analyzed cell lines (Extended Data. 1J), always show a significant enrichment in the SG common *core* (Fig. 1D). This may be due to their tendency to be longer than other gene categories (e.g. metabolic process, mitochondrion, endoplasmic reticulum and translation, Fig. 1G). Indeed, length was confirmed as a relevant feature for SG engagement in any cell type analyzed (Fig. 1E).

However, despite the presence of common *core* transcripts, the SG RNA content shows strong divergences in the different systems, with 37,8%, 40,4% and 52,7% of the SG transcriptomes differing in SK-N-BE, U2OS and SMMC-7721 granules, respectively. Interestingly, the SG transcriptome in neuroblastoma cells is enriched for neuronal categories such as “neuron to neuron synapse”, “neuronal cell body” and “axon part”, which are absent in the other systems (Fig. 1F, Extended Data Fig. 1K, L and M, Supplementary Table 2). These data indicate that, in our neural-like system, transcripts with neuronal functions have propensity to be included in SG, and this behavior correlates with their length compared to transcripts belonging to other categories (Fig. 1G). Furthermore, we observed that the length of the CDS and 3’UTR, and less of 5’UTR (Extended Data Fig. 1N), are crucial features for SG engagement.

However, length and RNA biotype are not the only discriminating features for recruitment into granules. In fact, when we analyzed the nucleotide composition, we observed that the SK-N-BE and SMMC-7721 datasets have a higher GC content than the U2OS set (Fig. 1E), suggesting that the inclusion of specific transcripts leads to different overall nucleotide composition and thus could be associated with different physical properties of SG. Noteworthy, these evidences are not dependent on the thresholds selected to define enriched RNAs since we observed the same results by stratifying the dataset according to the enrichment score (log_2_FC) in SG and comparing groups of equal numerosity (Extended Data Fig. 1O).

### ALS-related FUS mutation P525L affects SG dynamics and transcriptome composition

Mis-localization of mutant FUS in the cytoplasm and its accumulation in inclusions is known to be a central hallmark of ALS/FTD-associated mutations. Moreover, it is known that the mis-localized FUS co-localizes with other RBPs in SG as a consequence of oxidative stress, and that this process can lead to aggregation^33^. Despite this evidence, few efforts have been made to characterize the implications of mutant FUS localization on the physic-chemical features of SG and their RNA composition. To elucidate this aspect, we generated two stable SK-N-BE cell lines overexpressing under doxycycline control either the wild-type FUS protein (FUS^WT^) or a version of FUS carrying the P525L mutation on the NLS, which is associated to severe and juvenile ALS (FUS^P525L^)^34^. We induced FUS overexpression with 50 ng/mL doxycycline obtaining expression levels of ectopic FUS comparable to the endogenous protein (Extended Data Fig. 2A). We showed that the overexpressed FUS^P525L^ is physically recruited into SG, as demonstrated by biochemical purification (Extended Data Fig. 2B). We also verified the correct FUS localization and observed that, while overexpressed FUS^WT^ is completely localized in the nucleus, FUS^P525L^ is cytoplasmic and co-localizes upon stress with the SG core protein G3BP1 in specific foci (Extended Data Fig. 2C and 2D). Moreover, we compared the number of SG in both conditions. As reported in other systems^35^, we found an increased number of stress granules per cell in the presence of FUS^P525L^ (Extended Data Fig. 2E).

We then analyzed GFP-G3BP1 dynamics in FUS^WT^ and FUS^P525L^ cells, using fluorescence fluctuation spectroscopy with a SPAD array detector, to evaluate changes in the dynamics of condensation due to FUS mutation^36,37^. In agreement with previous observations, we found that in both cell lines G3BP1 movement decreases upon stress induction (Fig. 2A *-D*_app_, measures the speed of movement of individual molecules). The decrease of the diffusion coefficient upon stress induction reflects the internalization of G3BP1 inside SG. We then analyzed the diffusion times at different detection areas (integrating the fluorescence signal at different pixels of the SPAD array detector) to perform the so-called diffusion law. When the intercept of the linear regression of the diffusion times at different detection areas is bigger than zero, the sample has a domain-confined type of movement; when it is lower than zero, the sample has a meshwork-confined type of movement. The results of Fig. 2B show a more confined type of movement in presence of FUS^P525L^ compared to FUS^WT^, suggesting a less dynamic conformation for SG when the mutant FUS is included. Altogether, these data indicate that SG physiology is strongly affected by mutant FUS; in particular, the presence of FUS^P525L^ in SG induces an increased number of granules that are globally less fluid.

**Figure 2.**
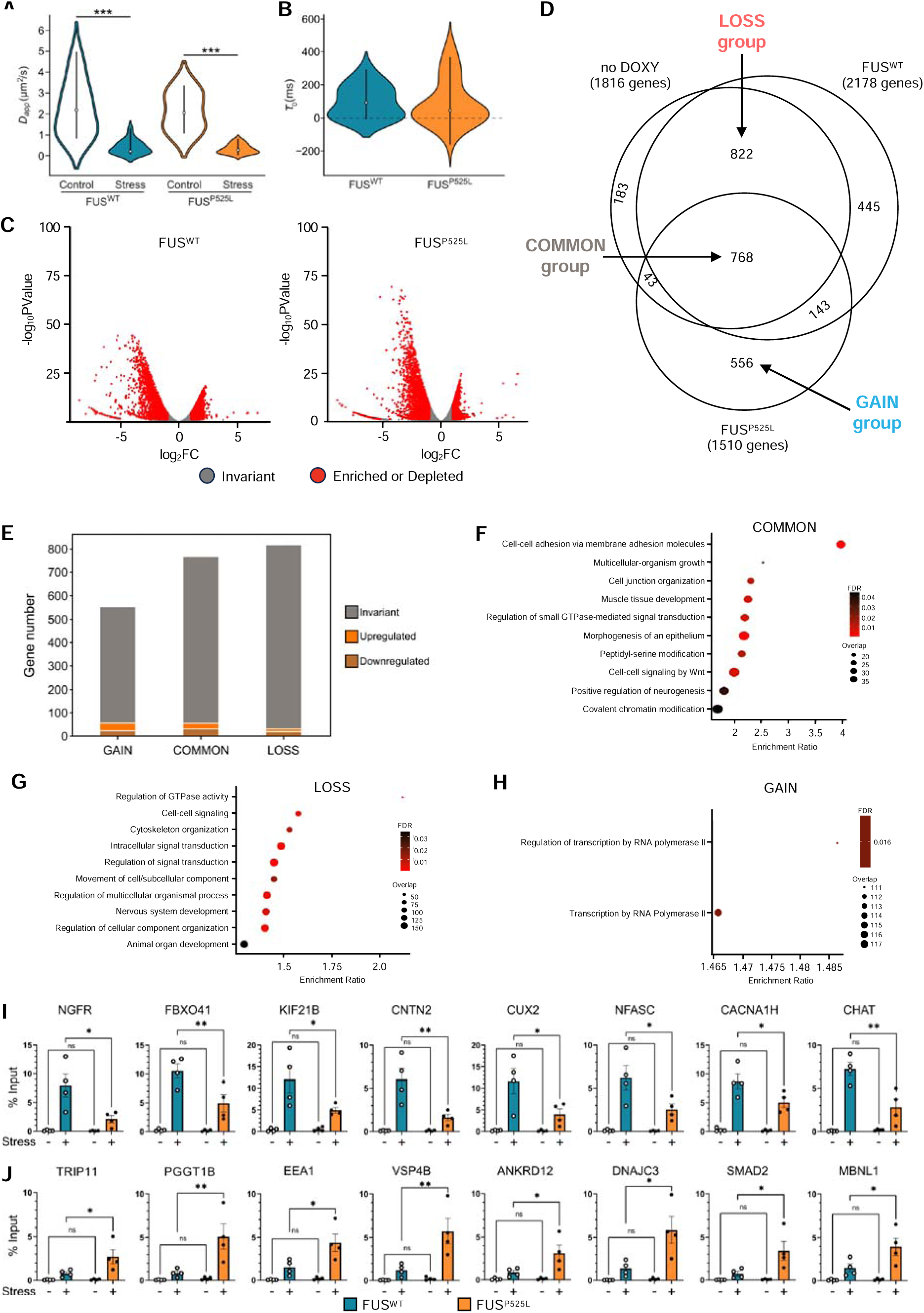
**A)** Violin plot showing spot-variation Fluorescence Correlation Spectroscopy measurements of the diffusion coefficient (*D*_app_) of GFP-G3BP1 in live SK-N-BE cells overexpressing FUS^WT^ and FUS^P525L^, both in control and stress conditions. *D*_app_ measures the speed of movement of individual molecules. **B)** Violin plot comparing the diffusion times (t_0_) of individual GFP-G3BP1 molecules within formed SG in FUS^WT^ and FUS^P525L^ conditions. The diffusion time t_0_ is obtained by integrating the fluorescence signal at different pixels of the SPAD array detector and defines the confinement of movement of the analyzed molecules. **C)** Volcano plot showing for each RNA detected in the RNA-seq experiment in FUS^WT^ (left panel) or FUS^P525L^ condition (right panel) the log2 fold change (SGenr/Input) and the -log10 p-value of SG enrichment. Significantly enriched or depleted RNAs (FDR < 0.05, |logFC| > 1) or invariant RNAs are indicated by red or gray dots, respectively. **D)** Venn diagrams showing the overlap between SG enriched RNAs in no DOXY, FUS^WT^ and FUS^P525L^ conditions. The amount of RNAs resulted enriched in SGs of each condition was indicated. Arrows indicate the fraction of RNAs indicated as FUS^P525L^ GAIN, COMMON and FUS^P525L^ LOSS. **E)** Stacked Barplot depicting the amount of deregulated RNAs in FUS P525L vs FUS WT INPUT comparison among the GAIN, COMMMON and LOSS groups. **F)** Dotplot depicting GO over-represented categories in the COMMON group. Only significant categories (FDR < 0.05) of Biological Process database were depicted. X-axis represents category enrichment score while y-axis reports the GO category description. Dot size represents the amount of RNAs in the analyzed group that overlap the category while color code (red) reports the significance of the enrichment. **G)** Dotplot depicting GO over-represented categories in the FUS^P525L^ LOSS group. Only significant categories (FDR < 0.05) of Biological Process database were depicted. X-axis represents category enrichment score while y-axis reports the GO category description. Dot size represents the amount of RNAs in the analyzed group that overlap the category while color code (red) reports the significance of the enrichment. **H)** Dotplot depicting GO over-represented categories in the FUS^P525L^ GAIN group. Only significant categories (FDR < 0.05) of Biological Process database were depicted. X-axis represents category enrichment score while y-axis reports the GO category description. Dot size represents the amount of RNAs in the analyzed group that overlap the category while color code (red) reports the significance of the enrichment. **I)** qRT-PCR analysis to validate the effective enrichment of RNAs selected from the LOSS group in samples of SG purification from SK-N-BE cells expressing FUS^WT^ (blue) or FUS^P525L^ (orange). Control and Stress condition were analyzed. Values are expressed as % of input, statistical significance was assessed through one-way ANOVA. n=4. **J)** qRT-PCR analysis to validate the effective enrichment of RNAs selected from the GAIN group in samples of SG purification from SK-N-BE cells expressing FUS^WT^ (blue) or FUS^P525L^ (orange). Control and Stress condition were analyzed. Values are expressed as % of input, statistical significance was assessed through one-way ANOVA. n=4.

Since we observed variations in SG number and dynamics, we decided to analyze whether the RNA content of these granules is altered in presence of FUS^P525L^. Therefore, we generated duplicated samples of purified SG and performed total RNA sequencing with matched total RNA inputs in condition of doxycycline induction of FUS^WT^ and FUS^P525L^. Leading Fold change analysis (Extended Data Fig. 2F) and dendrogram of relative correlation distances (Extended Data Fig. 2G) confirmed that samples show the expected clusterization in INP and SG-enr conditions and by FUS^WT^ or FUS^P525L^ backgrounds. Therefore, for each condition we compared the SG-enriched samples with matched input: in FUS^WT^ conditions we identified 2178 enriched (log_2_FC > 1 and FDR < 0.05) and 2990 depleted RNAs (log_2_FC < -1 and FDR < 0.05) (Fig. 2C, left panel); using the same thresholds, we observed 1510 SG enriched and 2752 depleted RNAs in FUS^P525L^ (Fig. 2C, right panel, Supplementary Table 1). Besides, we compared the SG transcriptome in the three datasets we generated: FUS no DOXY (without FUS induction), FUS^WT^ and FUS^P525L^. First, we observed that 87,5% of no DOXY SG RNAs are present also in FUS^WT^ SG, as expected from the mainly nuclear localization of the overexpressed WT protein. Conversely, investigating the impact of mutant FUS, we observed that 822 RNAs enriched in SG in both no DOXY and FUS^WT^ conditions are excluded from FUS^P525L^ SG (thus, we renamed this group of transcripts “LOSS”). Moreover, we also identified a consistent group of 556 RNAs that are specifically recruited in SG in presence of the mutation, but not in the other two conditions (group renamed as “GAIN”). Despite the strong differences observed, a set of 768 transcripts is maintained in all the three conditions and always engaged in SG (“COMMON” group) (Fig. 2D).

In order to check whether such alternations in SG composition could be due to diversities in the FUS^P525L^ and FUS^WT^ transcriptomes, we performed differential expression analysis in INP samples of FUS^P525L^ and FUS^WT^. The results indicate that the two conditions differ for only a small fraction of transcripts (Extended Data Fig. 2H - 244 among the downregulated transcripts and 274 in the upregulated ones) and that the contribution of such RNAs to the LOSS, COMMON and GAIN groups is negligible (Fig. 2E).

Moreover, using “Resampling analysis” (see Material and Methods), we confirmed that the observed alterations of SG transcriptome in presence of FUS^P525L^ is not dependent on the technical variability of RNA-Seq library sizes (Extended Data Fig. 2I), while “Enrichment convergence analysis” (see Material and Methods) confirmed that it is does not depend on the enrichment threshold (log_2_FC SG-enr vs INP) used to define the SG transcriptomes (Extended Data Fig. 2J).

To understand whether specific RNA categories are associated to the observed reorganization of SG transcriptome, we performed Gene Ontology terms enrichment analysis (Supplementary Table 2). RNAs that are enriched in all conditions (COMMON) include functions related to neural development, cell-cell adhesion and transcription regulators (Fig. 2F, and Extended Data Fig. 2K). Instead, in the LOSS group, we found RNAs related to cell signaling pathways, and cytoskeletal and neuronal components (Fig. 2G, and Extended Data Fig. 2L). Finally, in the GAIN group we found fewer categories mainly related to transcriptional regulation (Fig. 2H, and Extended Data Fig. 2M).

To biochemically validate the differential recruitment of individual RNAs into SG in the different conditions analyzed, we selected several transcripts enriched in the LOSS and GAIN groups, which were described to be linked to neurodegeneration or ALS (COMMON=311 genes, LOSS=321 genes and GAIN=172 genes; see Supplementary Table 2, tab “Literature survey” for references). Through qRT-PCR amplification on independent SG purification samples, we confirmed the strong difference in the ratio of recruitment of the selected RNAs due to the expression of mutant FUS (Fig. 2I and 2J). This is quite an interesting finding since both sets of transcripts can be equally vulnerable in the presence of mutant FUS: the GAIN group would suffer for the inability of FUS^P525L^ SG to be promptly dissolved upon stress release^38,39^ and would have a delayed translational recovery, while the LOSS one would not be protected from stress insults since excluded from SG. Altogether, these data suggest that the ALS-associated FUS^P525L^ mutation has a strong effect on SG dynamics and induces a dramatic reorganization of the SG transcriptome which includes components previously associated to neuronal degeneration.

### FUS^P525L^ affects SG transcriptome nucleotide composition

We then decided to investigate whether FUS mutation also affects RNA length, UTRs and CDS size and GC content of SG enriched RNAs that we previously identified. Thus, we considered how SG enrichment (log_2_FC) relates with those characterizing features.

As we observed for the no DOXY condition, length is a key feature to determine SG entry for both FUS^WT^ and FUS^P525L^ samples, with longer RNAs being more efficiently recruited (Fig. 3A). Moreover, we noticed that the length of CDS and 3’UTR of mRNAs is relevant for SG engagement in all the three conditions, while the contribution of the 5’UTR in this enrichment is less obvious. However, despite the length of RNAs and mRNAs regions being a common criterion for SG engagement, we could detect some relevant differences: RNAs in the FUS^P525L^ SG are globally longer (Extended Data Fig. 3A), and this increased size is mainly due to the length of the 3’UTRs. Indeed, conversely to what it was observed for the 3’UTR, both 5’UTR and CDS are slightly shorter in the FUS^P525L^ SG transcriptome (Extended Data Fig. 3B).

**Figure 3.**
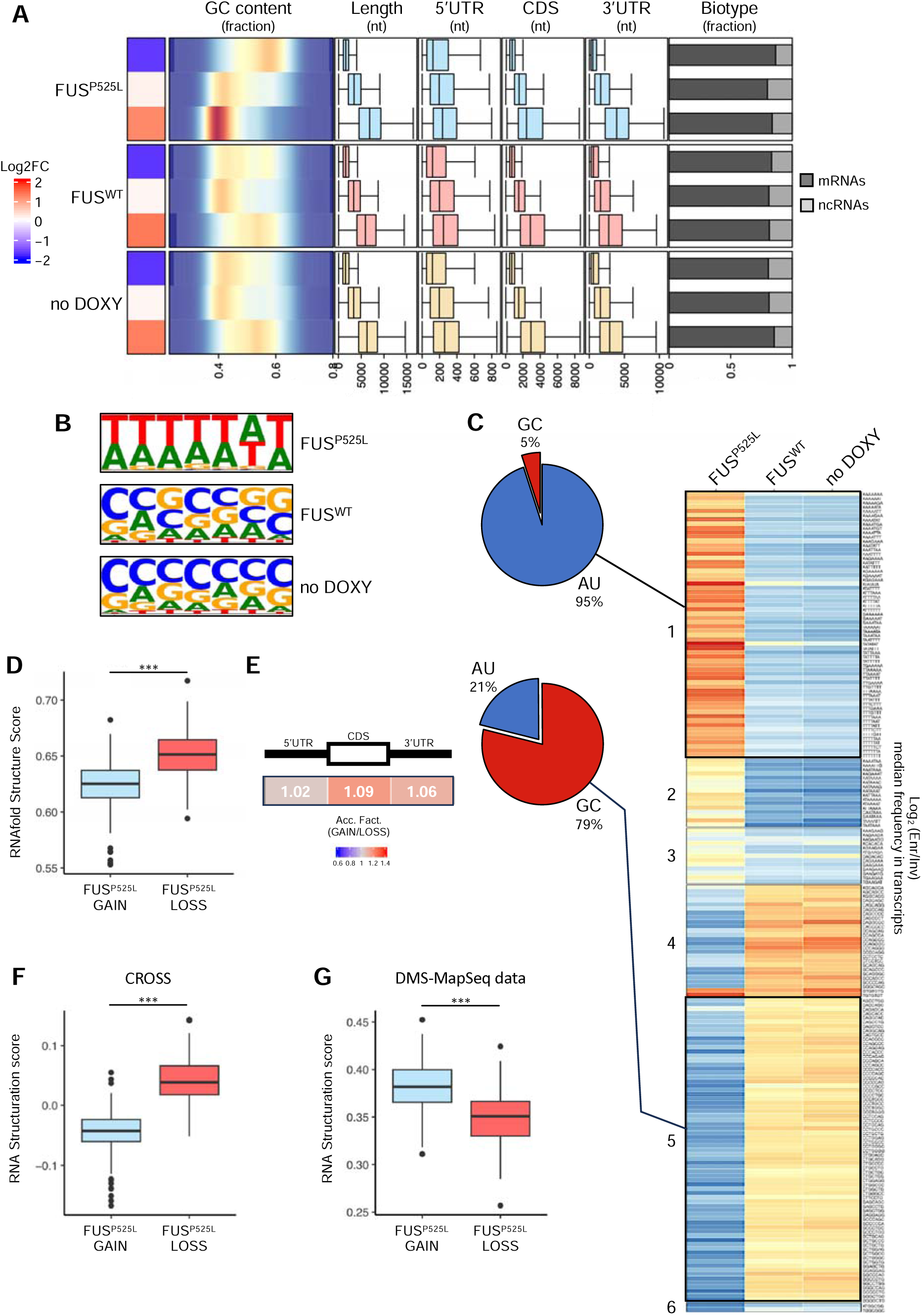
**A)** Heatmap depicting the association of RNA features with SG enrichment in FUS^P525L^, FUS^WT^ and no DOXY conditions. For the defined transcripts group («depleted», «invariant» and «enriched”) of each cell line the following characteristics were described: median log2FC (heatmap); GC content (density plot); RNA length (nt, boxplot); 5’ UTR, CDS and 3’UTR length (nt, boxplot); transcripts biotypes fraction (barplot). **B)** Sequence logo of the top 50 7-mers over-represented in SG enriched RNAs compared to SG depleted in no DOXY, FUS^WT^ and FUS^P525L^ conditions. **C)** Heatmap showing the clusters of 7mers frequency fold between SG enriched over invariant RNAs in no DOXY, FUS^WT^ and FUS^P525L^ conditions. For each 7mer, the log2 fold between average frequencies in groups was depicted. Only the union of the top 50 over-represented and 50 under-represented 7-mers in each condition was depicted. Pie charts represent the percentage of AU or GC nucleotides in the 7-mers enriched (cluster 1) and depleted (cluster 5) in FUS^P525L^ SG. **D)** Boxplot depicting the distribution of RNA structuration degree predicted using RNAfold algorithm between FUS^P525L^ GAIN and LOSS groups. For each RNA, the minimum free energy (MFE) structure was taken in consideration. The differences between groups were calculated using Mann-Whitney U test. The first sampling was reported. Size of sampling = 200 RNAs per group. **E)** Representative heatmap showing the fold between average accessibility of FUS^P525L^ GAIN over LOSS groups. The differences between groups were calculated using Mann-Whitney U test. Only significant differences were depicted (P-value < 0.05). The first sampling was reported. Size of sampling = 200 RNAs per group. **F and G)** Boxplot depicting the distribution of RNA structuration score predicted by CROSS algorithm (F) and RNA average accessibility calculated using DMS-Map-Seq data (G) between FUS^P525L^ GAIN and LOSS groups. The differences between groups were calculated using Mann-Whitney U test. The first sampling was reported. Size of sampling = 200 RNAs per group for CROSS (F) and 50 RNAs per group for DMS-Map-Seq (G).

Furthermore, we also analyzed the fraction of protein coding transcripts and we observed that most of the RNAs identified as enriched were mRNAs, with comparable percentages in all the three groups (80% in FUS^WT^ to 85% FUS^P525L^ and 85% no DOXY) (Fig. 3A).

However, we observed the strongest difference in the GC content of the compared transcriptomes, with a consistent reduction of the GC content in the FUS^P525L^ condition compared to no DOXY and FUS^WT^ conditions (Fig 3A and Extended Data Fig. 3C). Indeed, analyzing the GC content in no DOXY and FUS^WT^ conditions, we observed a low correlation degree with SG enrichment (Rho=0.065 no DOXY and Rho=0.04 FUS^WT^, in Extended Data Fig. 3D), but when we focused on the FUS^P525L^ condition we strikingly observed a strong anti-correlation (Rho=-0.3, Extended Data Fig. 3D). This evidence not only highlights that SG enriched RNAs in FUS^P525L^ condition have a greater AU content (Fig 3A) compared to the others, but also that this enrichment is a feature associated with RNA entry in SG only in the presence of mutant FUS.

Noteworthy, this evidence is not dependent on the thresholds selected to define enriched RNAs. Indeed, we got the same results by stratifying the dataset according to the enrichments score in SG (Extended Data Fig. 3E).

To further investigate the relationship between sequence composition and SG recruitment, we asked whether specific subsequences are particularly represented in SG enriched transcripts. To highlight this aspect, we performed a k-mer (k = 7) enrichment analysis. Analyzing the top 50 k-mers over-represented in SG RNAs, we found that transcripts enriched in both FUS^WT^ and no DOXY conditions are characterized by a higher content of GC-rich k-mers, while the mutant FUS condition preferentially recruits RNAs containing AU-rich k-mers (Fig. 3B). Indeed, K-means clustering analysis of subsequence enrichment in SG identified two defined clusters of k-mers (cluster1 and cluster 5), that have divergent patterns of enrichment: the cluster 1, specifically enriched in FUS^P525L^, displayed AU-rich sequences, while the cluster 5 that is specifically enriched in no DOXY and FUS^WT^ conditions displayed GC-rich ones (Fig. 3C).

To better understand the relationship between different RNA composition and the phenotype of reduced SG dynamics observed in FUS^P525L^, we decided to deeper investigate the properties of the LOSS and GAIN transcripts, which represent the strong changes in transcriptome architecture. To do that, we compared the RNA secondary structures propensity of mRNAs from the two groups. In order to have a more reliable comparison, we sampled an equal number of RNAs from the two groups; moreover, since mRNA functional regions (5’UTR, CDS and 3’UTR) have different structural and interaction properties, we compared mRNAs with similar 5’UTR, CDS and 3’UTR length (Mann Whitney U-Test, p-value > 0.05, Extended Data Fig. 3F, G and H). We then applied the RNAfold algorithm^40^ to predict the minimum free energy (MFE) RNA secondary structure of each transcript, calculating the structuration score as the number of paired nucleotides over the length of the full transcripts or regions of them. Interestingly, we found that RNAs in the GAIN group were globally less structured than the LOSS group (Fig. 3D and Extended Data Fig. 3I). Moreover, comparing accessibility (defined as the fraction of un-paired nucleotides over the length of the full transcripts or regions of them) in each mRNA region, we found that CDS displays the stronger contribution to this feature, with 9% increased accessibility in FUS^P525L^ GAIN compared to the FUS^P525L^ LOSS group (Fig. 3E).

Moreover, to corroborate this evidence with other approaches, we used CROSS algorithm^41^ that is an RNA secondary structure predictor based on a machine learning approach, with parameters trained on RNA structures derived by PARS datasets. We observed a higher prediction score for LOSS RNAs compared to the GAIN group (Fig. 3F and Extended Data Fig. 3J), highlighting that the RNAs that lose their enrichment in SG in presence of mutant FUS are more structured. Finally, in accordance with the predictions, also RNA accessibility data derived from DMS-Map-Seq methodology^42^ confirmed less single stranded regions in the RNAs of LOSS group compared to the GAIN ones (Fig. 3G and Extended Data Fig. 3K). Altogether, these data suggest that FUS^P525L^-mediated alteration of SG has a strong impact on the global GC content of the granules, and that the RNAs specifically lost in this condition are more structured, while the recruited ones are less prone to generate secondary structures.

### FUS^P525L^ mutation alters the binding properties of the SG proteome

In order to investigate the causal links between the presence of mutant FUS and the observed alterations in SG transcriptome, we focused on its RNA interactome. Indeed, since mutant FUS is almost completely localized in SG and directly binds RNAs, we expected an enrichment of FUS interacting transcripts in FUS^P525L^ SG compared to the WT condition.

We investigated this possibility by taking advantage of the endogenous mutant FUS PAR-CLIP experiment performed in iPSC derived motor-neurons^43^. Remarkably, even though FUS establishes interactions with a wide fraction of the transcriptome, we found a significant, yet subtle increase of about 10% of interacting RNAs when comparing FUS^P525L^ binders between the GAIN and LOSS groups, suggesting that FUS direct binding is not sufficient to explain the differential recruitment of transcripts in the SG of WT versus mutant conditions (Fig. 4A). Moreover, we found a similar SG recruitment rate of FUS RNA interactors in the FUS^P525L^, no DOXY and FUS^WT^ conditions, when comparing classes of RNAs with lower or higher rate of enrichment (log_2_FC) into SG (Extended Data Fig. 4A).

**Figure 4.**
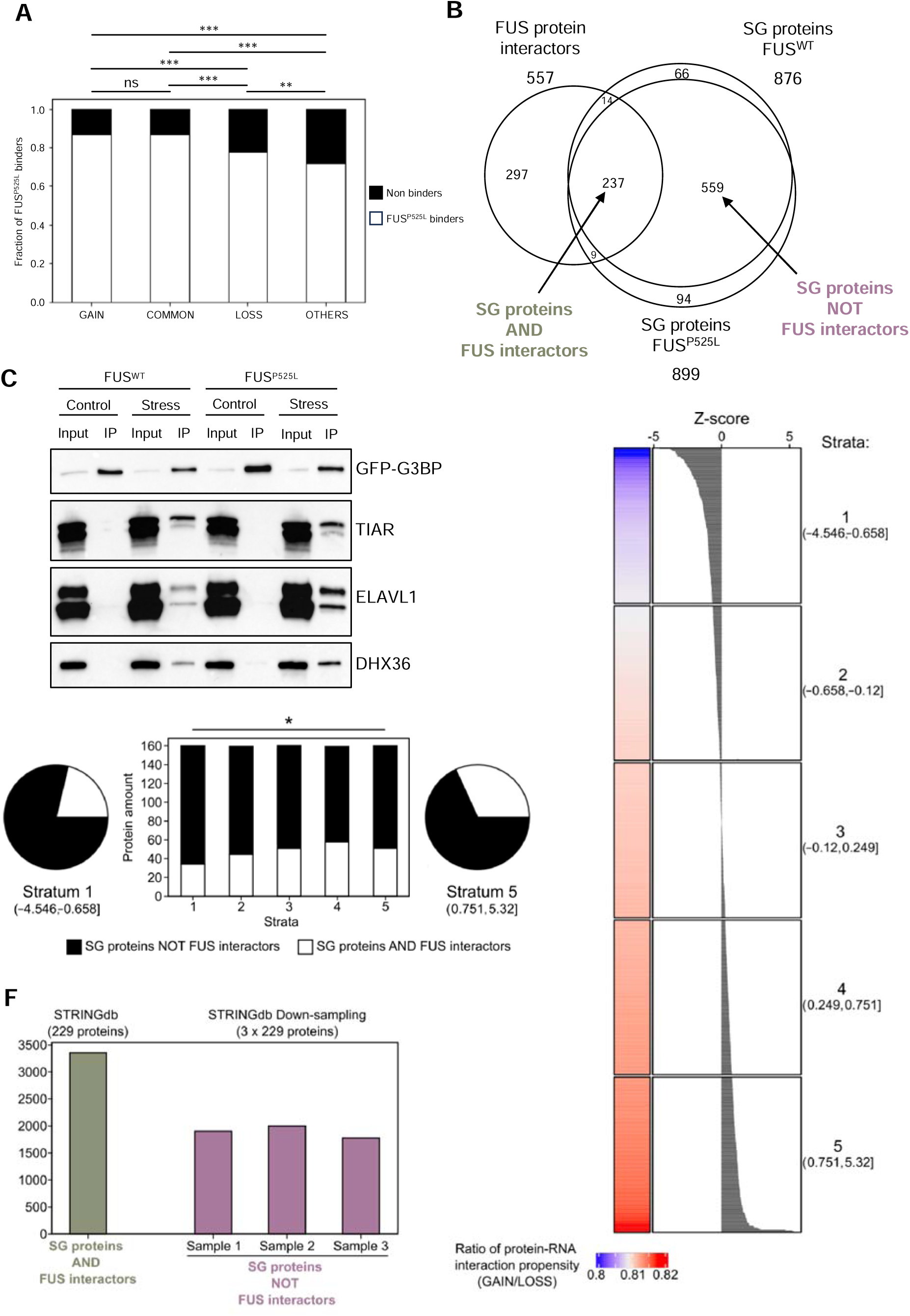
**A)** Barplot depicting the enrichment of endogenous FUS^P525L^ PAR-CLIP interactors retrieved from iPSC derived motor-neurons (MNs) in GAIN and COMMON compared to LOSS groups, and the remaining fraction of the transcriptome. Only RNAs detected in both systems (SK-N-BE and MNs) were taken in consideration. Black color refers to the fraction of RNAs not bound by FUS, while white color refers to FUS^P525L^ RNA interactors. The significance of enrichment was calculated with Fisher’s exact test (two-tails), after multiple-test correction (FDR). **B)** Venn diagram depicting the overlap among SG proteomes in FUS^WT^ and FUS^P525L^ conditions and FUS protein interactors. Arrows indicate the fraction of proteins defined as SG proteins AND FUS interactors or SG proteins NOT FUS interactors. **C)** Representative Western Blot analysis of GFP-G3BP1, TIAR, ELAVL1 and DHX36 in samples of SG purification, in both FUS^WT^ and FUS^P525L^ conditions. The immunoprecipitation (IP) sample was compared with 5% of total protein Input (Input). Control and Stress conditions were compared. n=3. **D)** Heatmap depicting the SG proteome ranked by interaction propensity for GAIN over the LOSS group. Heatmap colors from blue to red report the ratio of SG proteome average interaction propensity (RNAact, catRAPID) of GAIN over LOSS groups, the barplot on the left reports the Z-score of these ratios. The SG proteome was stratified in 5 groups with first group (bottom) that displayed the highest interaction propensity for the GAIN group, while the last group (top) displayed the highest interaction propensity for the LOSS group. **E)** Pie charts depicting the fraction of SG proteins AND FUS interactors and SG proteins NOT FUS interactors in the first (left) and last group (right). Bar plot (middle) displays the indicated fraction in every stratification group. Black color refers to SG proteins NOT FUS interactors while white color refers to SG proteins AND FUS interactors. For GAIN and LOSS comparison the matched length sampling were used. The first sampling was depicted. Significance of the enrichment of SG proteins AND FUS interactors in the first group compared to the last was assessed with Fisher’s exact test. These data refer to the analysis on RNA-protein interaction made with catRAPID algorithm. **F)** Barplot depicting the enrichment of protein-protein physical interactions (PPI) in the SG proteins AND FUS interactors compared to the three samplings performed from the SG protein NOT FUS interactors group.

Therefore, we searched for other possible contributions to this remodeling of SG-recruited RNAs, hypothesizing that the presence of mutant FUS could alter the proteome, thus indirectly impacting on the SG transcriptome. For this reason, aiming to characterize the protein content of granules in FUS^WT^ and FUS^P525L^ conditions, we performed SG purification followed by mass spectrometry, detecting 876 and 899 proteins in the two conditions, respectively (Supplementary Table 3). In order to assess the proper purification and detection of the SG proteome, we performed a GO enrichment analysis on the proteins identified in FUS^WT^ condition; since the overexpressed FUS^WT^ is almost exclusively nuclear, these samples could be used as reference control. As expected from already described SG proteomes^17,44^, we found among the most enriched categories those related to translation, RNA processing and RNA binding (Extended Data Fig. 4B, Supplementary Table 3).

Moreover, we compared the SG proteome in FUS^WT^ condition with other published SG proteome datasets from HEK293T cells^45^ and U2OS cells^17^, observing that a consistent portion of the SG localized proteins (77 common *core* proteins) are shared by all the three systems (Extended Data Fig. 4C). Among the common *core* proteins, we found known SG components such as G3BP2 and CAPRIN1, helicases such as DHX36, RNA processing factors like XRN1 and CNOT1, and also RNA binding proteins such as STAU2, PUM1 and EWSR1 (Extended Data Fig. 4D). Despite the common *core*, we also found proteins identified only in our dataset and not in the others (Extended Data Fig. 4C), supporting the idea that the common SG *core* is accompanied by another set of proteins specific for the system under analysis^46^.

Therefore, we studied the impact of FUS^P525L^ on the SG proteome by comparing the identified proteins from mass spectrometry data in FUS^WT^ or FUS^P525L^ conditions. Conversely to what we observed for SG transcriptome, we found strong similarities between the proteomes in both conditions. Indeed, among the 876 proteins detected in FUS^WT^ SG and the 899 proteins in FUS^P525L^ SG, 796 are shared by the two conditions (90,8% of FUS^WT^ and 89% of FUS^P525L^ respectively, Fig. 4B), suggesting that the presence of mutant FUS does not cause a dramatic inclusion of new protein species in the SG. However, despite the qualitative similarity in SG protein contents, the increase of mutant FUS due to its cytoplasmic de-localization itself could have an impact on the quantitative recruitment of SG-associated proteins. Thus, in order to better understand the relationship between FUS protein interactors and the SG proteome, we took advantage of 5 published datasets of FUS protein interactors related to its soluble^47–49^, condensed^50^ or aggregated form^45^. The latter dataset refers to a mis-localized and aggregated form of FUS carrying the R522G mutation, and displayed the highest amount of FUS protein interactors (488 proteins). Interestingly, across all datasets we found that a large fraction of FUS interactors is present in SG in both WT and mutant conditions, starting from a minimum of 40% up to 85% of overlap across the five interactomes analyzed (Fig 4B, Extended Data Fig. 4E), meaning that a high portion of FUS protein interactors are already SG constituents. This evidence supports the hypothesis that mutant FUS is able to interact with a consistent set of proteins engaged in SG.

Interestingly, known FUS interactors, such as DHX36 (also known as RHAU) and ELAVL1 (HuR), widely studied for their ability to bind AU-rich sequences^51,52^, were found to be more recruited in SG in the presence of FUS^P525L^, as demonstrated by Western Blot in comparison with another unperturbed SG marker (TIAR) (Fig. 4C). Since the expression of these factors is comparable in the Input fractions of control and mutant conditions (Extended Data Fig. 4F), their higher levels in FUS^P525L^ SG suggest that they can cooperate with FUS for the strong remodeling of SG transcriptome.

To check whether RNAs characterizing the FUS^P525L^ GAIN group (those recruited in SG specifically upon FUS mutation) are preferentially bound by FUS protein interactors, we took advantage of RNAact database of protein-RNA interaction predictions based on catRAPID algorithm^53^. To do that, we ranked the SG proteome for the preferential binding with the GAIN RNA group compared to the LOSS one (Fig. 4D). Interestingly, among the SG proteins that are more prone to interact with the GAIN RNAs, we found an enrichment of proteins that interact with FUS (Fig. 4E, Extended Data Fig. 4G). We noticed a similar trend of enrichment of FUS interactors as preferential binders of the GAIN RNAs over the LOSS ones even when we performed the same analysis using experimental data of RBP-RNA interactions from POSTAR3 database^54^ (Extended Data Fig. 4H). This evidence suggests that FUS^P525L^ mis-localization and increased cytoplasmic concentration favors the SG recruitment of both its RNA and protein interactors; moreover, the increased engagement of FUS protein interactors into SG could be responsible for the alteration of SG transcriptome by driving the inclusion of the RNAs of the FUS^P525L^ GAIN group.

We also investigated the impact of the inclusion of mutant FUS into SG on the interaction propensity of the SG proteome. Thus, we compared the protein-protein physical interactions (PPI, STRING database) that are expected to occur in the group of SG FUS interactors (named “SG proteins AND FUS interactors”) and in the remaining fraction of SG proteins (named “SG proteins NOT FUS interactors”). Since these two groups have different amounts of proteins (237 and 559, respectively, Fig. 4B), we performed three down-sampling of “SG proteins NOT FUS interactors” group and compared the PPI of “SG proteins AND FUS interactors” with those of the down-sampled groups to obtain a more reliable comparison (Extended Data Fig. 4I and 4J). Interestingly, the SG proteins that don’t interact with FUS show around 40% less PPIs than FUS interactors (Fig. 4F and Extended Data Fig. 4K). This evidence indicates that mutant FUS localized in SG could affect the hub of protein-protein interactions of a huge set of SG components.

Taken together, our data suggest that the recruitment in SG of mutant FUS produces alteration of the transcriptome both through engagement of FUS direct RNA interactors, as well as by the recruitment of FUS protein interactors which in turn could convey in SG their bound RNAs; moreover, we suggest that the inclusion of FUS^P525L^ protein partners could reshape the protein-protein interaction landscape inside the granules.

## DISCUSSION

In this study, we addressed how ALS-associated mutation of the RBP FUS affect SG molecular composition in a neural-like context.

Combining granule purification with high throughput RNA sequencing, we characterized the composition of SG RNAs in a human neural-like system (SK-N-BE, neuroblastoma), highlighting the recruitment of neuronal-specific RNAs. Also in our conditions, length is the main feature of the SG-enriched RNAs, in agreement with the description of SG RNA composition in other biological systems^31,32^. Since longer RNA molecules usually harbor an increased number of possible interaction sites with both other RNAs and proteins, this feature emphasizes the role of multiple non-specific RNA interactions in the phase separation process^8^. Moreover, we described the GC content as a diversifying feature of SG RNA in different cell types: the high GC content of neuronal SG could give them peculiar properties in term of RNA-protein interaction networks^55^.

Moreover, using SK-N-BE cells expressing FUS^WT^ and FUS^P525L^, we were able to reveal important physical alterations of SG condensates in which mutant FUS is embedded. Indeed, recruitment of FUS^P525L^ to SG increases their number and alters their molecular dynamics, leading to more aggregated SG with reduced physical mobility.

Furthermore, RNA-seq analyses revealed that FUS^P525L^ altered the RNA composition of neural SG, resulting in the inclusion of a new set of transcripts (GAIN group) and the exclusion of others (LOSS group), if compared to FUS^WT^. Among the re-localized RNAs we found several transcripts described to be linked to neurodegenerative phenotypes and ALS. This is of interest since both GAIN and LOSS transcripts could be both affected by the presence of mutant FUS in the cytoplasm. On one hand, the GAIN group RNAs are entrapped in granules that are more resistant to dissolution, and would suffer for the inability of FUS^P525L^ SG to be promptly dissolved upon stress release^38,39^ and to rapidly recover translational. On the other hand, the LOSS group RNAs would not be protected from stress-related damage since excluded from SG^56^.

FUS mis-localization was also shown to strongly alter the GC-content of recruited RNAs, shifting towards AU-rich sequences; the observed alteration in nucleotide composition implies a different structuration propensity of the SG transcriptome. Indeed, lost RNAs result more likely to form secondary structures compared to the gained ones. This GC divergence is important to take into consideration in pathological contexts, since the RNA nucleotide composition is a crucial property for interaction with other RNAs and proteins^57^ and its alteration could be at the base of the susceptibility to the irreversible aggregation process of these granules.

Finally, analyzing the FUS^P525L^ RNA interactome^43^, we observed an enrichment of its direct RNA interactors, suggesting that a portion of transcripts is already associated with the mis-localized protein in the absence of stress, and that the formation of SG causes their engagement in the condensates. However, the extent of this enrichment cannot fully explain the transcriptome alteration observed in the granule, which is in turn justified by the engagement of other RNA binding proteins, partners of FUS, that could indirectly contribute to the observed RNA compositional changes. Despite the fact that we did not observe a strong qualitative difference in the proteome components of SG in presence of mutant FUS, we found that some protein interactors of FUS are more engaged into SG and that they could convey their RNA targets into these granules. Therefore, the modification of the SG transcriptome results from a double contribution of mutant FUS, which directly binds a portion of the SG recruited RNAs, but also drives the engagement of a subset of protein partners and, indirectly, of their interacting RNAs.

Furthermore, we have also observed that the FUS-interacting protein components of SG exhibit a significantly higher potential to form protein-protein physical interactions; this characteristic has been previously documented for the aggregated state of FUS^R522G^ interactors^45^. Moreover, the physical interactions considered for this analysis include those involved in the formation of protein complexes (STRING database) and extend beyond contacts simply involving the low-complexity domains (LCDs) of proteins. This observation further highlights the importance of protein-protein interactions in the formation of aggregates and less dynamic SG.

While it has been suggested that RNA itself possesses aggregating potential^58^, and undoubtedly contributes to the phenomena of phase separation, recent evidence supports the idea that RNA can act as a buffer in neurotoxic aggregation processes. This has been observed for TDP-43 and FUS, where the role of the RNA-binding domain (RRM) and its interaction with RNA is vital in constraining LCD-driven aggregation^59^. Furthermore, this finding aligns with a key paradigm of neurotoxic aggregation as a cytoplasmic event: condensation is influenced by the RNA-to-protein ratio, and in the nucleus this is limited by the abundance of newly transcribed RNA^60^.

From our observations, we suggest that the altered dynamics of SG in FUS^P525L^ conditions and the increased condensate aggregation state are associated with interesting molecular features: i) re-localization of relevant neuronal transcripts in SG, ii) reduced architectural capacity of RNAs, involving a decreased propensity to form secondary structures, and iii) increase in protein-protein interactions. In conclusion, our results shed new light on the importance of RNA features in the aggregation process involving SG. This not only suggests that specific and distinct RNA characteristics are linked to pathological granules and aggregation processes, but also introduces intriguing possibilities for understanding how these phenomena can impact neural physiology and be causatively linked to neurodegeneration.

## Supporting information

Supplemental Table 1 - RNAseq

Supplemental Table 2 - GO analysis

Supplemental Table 3 - Mass Spec

## AUTHOR CONTRIBUTION

I.B., D.M. and A.S. conceptualized the project. I.B., G.V and A.A. supervised and coordinated the experiments. D.M. set up the experimental system. D.M., F.C., and L.S.M. performed molecular biology experiments. A.S. designed and carried out all the bioinformatic analyses. E.V. performed and analyzed IF experiments. G.d.T. and A.G. analyzed results and discussed data. F.C., E.P. and S.Z. performed the FCS experiments. N.L. performed mass spectrometry. I.B., D.M. and A.S. wrote the original manuscript with contribution of all the authors.

## ACKNOWLEDGMENTS

We thank Prof. Alberto Diaspro and Dr. Paolo Bianchini (Nanoscopy & NIC@IIT, Istituto Italiano di Tecnologia), Dr. Michele Oneto (Nikon Imaging Center) and Marco Scotto (Molecular Microscopy and Spectroscopy, Istituto Italiano di Tecnologia) for experimental support in confocal and SIM microscopy. We are also grateful to Dr. Diego Vozzi and the Genomic Facility of CHT for support in RNA sequencing experiments. A special thank goes to the technical and administrative staff of IIT, and to all the members of the RNA Initiative@IIT community for thought-provoking discussions. We also wish to thank Angelo d’Angelo for fruitful discussion regarding bioinformatic analyses. The project was partially supported by grants from ERC-2019-SyG 855923-ASTRA, AIRC IG 2019 Id. 23053, PRIN 2017 2017P352Z4 and “National Center for Gene Therapy and Drugbased on RNA Technology” NextGenerationEU PNRR MUR (CN00000041), together with intramural IIT funding.

## DATA AVAILABILITY

All software, links to websites or tools used for this work are referred to in the Materials and Methods section or in the figure legends. Additional dedicated scripts developed for this work are available upon request.

## MATERIAL AND METHODS

### Cell culture, maintenance, and treatment

SK-N-BE were cultured in adherence in RPMI-1640 supplemented with 10% FBS, 1 mM Sodium Pyruvate, 1x GlutaMAX^TM^, 100 U/mL Penicillin and 100 µg/mL Streptomycin (all reagents were purchased from Thermo Fisher Scientific). Cells were frozen in SK medium supplemented with 20% FBS and 10% DMSO (Sigma Aldrich), and quickly thawed when needed in a 37°C water bath. To induce FUS overexpression, cells were incubated with 50 ng/mL Doxycycline hyclate (Sigma Aldrich) in SK medium for 24 hours. To induce Stress Granules formation, cells were incubated with 0,5 mM Sodium Arsenite (Sigma Aldrich) at 37°C for 1 hour; the effective condensation of GFP-G3BP1 was inspected under a fluorescence microscope before cell collection.

### Cloning and SK-N-BE stable cell line generation

The plasmid containing an N-terminal GFP-tagged form of human G3BP1 was a gentle gift from Roy Parker’s Lab (University of Colorado Boulder). The GFP-G3BP1 sequence was subcloned in the ePB-BSD-EIF1a vector^61^; briefly, GFP-G3BP1 was PCR-amplified with CloneAmp HiFi PCR Premix (TakaraBio) and cloned in the acceptor plasmid linearized with XbaI and NotI restriction enzymes (Thermo Fisher Scientific) using the InFusion Cloning kit following manufacturer protocol (TakaraBio); the NotI site was not maintained to reduce the Tm of the cloning primers. Primer sequences are listed in Supplementary Table 4.

Stable cell lines were generated taking advantage of the transposon-based PiggyBac technology. A PiggyBac HyPB transposase expression plasmid^62^ was co-transfected with the ePB-EIF1a-GFP-G3BP1 vector with a 1:10 ratio; cells were selected in 10 μg/ml of blasticidin S (Thermo Fisher Scientific) for 7 days and then tested for consistent expression of the fluorescent protein. SK-N-BE cell lines expressing FUS^WT^ and FUS^P525L^ under a doxycycline-inducible promoter were already present in the lab^63^.

### RNA extraction and analysis

RNA extraction was performed with the Direct-zol Miniprep RNA Purification Kit (Zymo Research) with on-column DNase treatment, according to the manufacturer’s instructions, with elution in 25 µL of RNase-free water (50 µL for total RNA Input samples). Total RNA was retro-transcribed with PrimeScript™ RT Reagent Kit (TakaraBio) according to the manufacturer’s instructions. For qPCR analysis, the PowerUp SYBR Green Master Mix (Thermo Fisher Scientific), following the manufacturer’s protocol, in 15 µL reactions, on a QuantStudio5 qPCR system (Thermo Fisher Scientific) with standard cycling conditions. For SG purification experiments, data were expressed as % of input: briefly, the Ct of Input samples for each target were adjusted to account for the dilution by subtracting the log2 of the dilution factor. Then, the % of input was calculated as 2^ (Adjusted input Ct - IP Ct) for each target. The sequences of primers used for qRT-PCR are listed in Supplementary Table 4.

### Literature survey

Literature survey analysis was performed using Entrez.esearch function of Biopython python library (https://academic.oup.com/bioinformatics/article/25/11/1422/330687). Bibliographic results indicated by PUBMED IDs in Supplementary Table 2 were obtained by searching on PubMed database the combination of Key words ("Amyotrophic Lateral Sclerosis" and "Neurodegeneration") with the genes belonging to COMMON, LOSS and GAIN groups linked by "AND" boolean logic proposition. We considered ALS linked genes also ANKRD12 and VSP4B, although not resulting from the described literature investigation. Indeed, ANKRD12 was described as down-regulated in murine spinal cord of TDP-43 N390D/+ mutants^64^, while VSP4B is transcriptionally regulated by TDP-43 and its derepression upon TDP-43 down-regulation leads to dendritic loss^65,66^.

### Protein extraction and Western Blot analysis

Protein extracts were obtained using RIPA buffer, supplied with 1× Complete Protease Inhibitor Cocktail (Merck). Protein concentration was assessed using BCA protein assay (Thermo Fisher Scientific). Protein electrophoresis was performed using 4–15% Mini-PROTEAN TGX Precast acrylamide gel (Bio-Rad), and proteins were transferred to 0,45 µm nitrocellulose membrane, using the TransBlot Turbo System (Bio-Rad), according to the manufacturer’s instructions. Membranes were stained with Ponceau, blocked with 5% non-fat dry milk (Sigma Aldrich) for 1 h and incubated overnight at 4°C with the following primary antibodies: anti-GFP (Thermo Fisher Scientific, A-11122, 1:1000), anti-TIAR (BD Transduction, 610352, 1:1000), anti-FUS (SantaCruz Biotech, sc-47711, 1:2000), anti-GAPDH-HRP (SantaCruz Biotech, sc-47724, 1:1000), anti-FLAG (Merck, F3165, 1:1000), anti-ELAVL1 (SantaCruz Biotech, sc-5261, 1:1000), anti-DHX36 (Proteintech, 13159-1-AP, 1:500). After secondary antibody incubation for 1 hour at room temperature, protein detection was carried out with Immobilon Crescendo Western HRP substrate (Merck) using ChemiDoc™ MP System and images were analyzed using Image Lab™ Software (Bio-Rad).

### Fluorescent microscopy analysis

Cells were plated on glass coverslips and treated as previously described. Samples were fixed in 4% paraformaldehyde for 10 min at room temperature (RT), followed by 3 washes in complete PBS. Permeabilization and blocking were performed by incubating the coverslips in PBS supplemented with 2% BSA, 0,2% TRITON X-100 for 30 min at RT. Samples were then incubated with primary antibody diluted in PBS with 1% BSA for 1 hour at RT, followed by 3 washes for 5 min in PBS. Fluorophore-conjugated secondary antibodies (Thermo Fisher Scientific) were diluted 1:300 in PBS with 1% goat/donkey serum and incubated for 45 min at RT. After 3 PBS washes, nuclei were stained with 4’,6-Diamidino-2-Phenylindole (DAPI, Sigma; diluted 1ug/ml in PBS) for 3 min at RT and coverslips were mounted with Prolong Diamond Mounting Media (Thermo Fisher Scientific, P-36961) on microscope slides. Fixed samples were imaged on a Nikon Instrument A1 Confocal Laser Microscope equipped with 60x and 20x objectives. Confocal images were collected with NIS-Elements AR software (Nikon): ND Acquisition module was used for multipoint acquisition of Z-stack images (150-175nm Z-spacing) of 4 um thickness.

Fluorescence Correlation Spectroscopy (FCS) measurements were performed as described in^36,37^. Cells were plated and FUS expression was induced as previously described; before starting the acquisition, culture medium was replaced with FluoroBriteTM DMEM (Thermo Fisher Scientific) supplemented with 10% FBS, and sodium arsenite was added at a final concentration of 0.5 mM for the “stressed” samples. Acquisition of GFP/RFP fluorescence signals was obtained in a custom-built laser scanning microscope equipped with a single-photon array detector from the Molecular Microscopy and Spectroscopy division of the Italian Institute of Technology in Genoa. A variant of conventional FCS (spot-variation FCS) was employed to quantify the diffusion mode of GFP-G3BP1. In spot-variation FCS, multiple diffusion times are measured at different detection volume sizes. Thanks to the spatial information granted by the single-photon array detector^36,67,68^, spot-variation FCS can be implemented within single measurements, by integrating off-line the signal coming from different subsets of the pixel detector, such as the central pixel only, the sum of the inner 3×3 pixels, and the sum of the entire 5×5 pixels. The diffusion modes are evaluated by fitting a linear function (diffusion law) to the diffusion times against the area of the detection volumes^69,70^. When the intercept of the linear function is zero, the molecules freely diffuse in the detection area; when the value is higher than zero, the molecules are isolated in domains. Conversely, when it is lower than zero, the molecules are constrained inside a meshwork.

### SG purification

Stress granules purification for the RNA sequencing experiments was performed according to^30^, with minor adjustments. After lysis, the protein concentration of samples was measured through BCA assay (Thermo Fisher Scientific); 3 mg of lysate were diluted in 1 mL of SG Lysis buffer and used for the purification, saving 5% as Total RNA input (stored in TRIZOL L/S, Thermo Fisher Scientific). SG were pelleted as in^30^, while the immunoprecipitation step was performed by incubation with antibody-coupled beads. For each sample, 50 µL of DEPC-treated Protein G Dynabeads (Thermo Fisher Scientific) were incubated with 12.5 µg of anti-GFP antibody in rotation at 4°C for 1 hour, then washed thrice in 1 mL of SG Lysis Buffer and finally resuspended in 150 µL of SG Lysis Buffer. Prepared beads were added to the precleared SG-enriched fraction and incubated in rotation at 4°C for 2 hours. Washes and beads elution were performed as in^31^. To isolate SG for qRT-PCR analysis, mass spectrometry and Western Blot validation, Dynabeads were replaced with 25 µL/sample of GFP-Trap beads (ChromoTek), which are covalently bound to a highly efficient alpaca-raised nanobody against the tag. For qRT-PCR, beads were washed and RNA was eluted as in^30^. To isolate proteins for mass spectrometry and WB analysis, after capture the beads were washed twice with 1 mL Wash Buffer (10 mM TRIS-HCl pH 7.5, 150 mM NaCl, 0.05% Igepal, 0.5 mM EDTA) in rotation for 10 minutes at 4°C, then washed twice with 1 mL High Salt Wash Buffer (10 mM TRIS-HCl pH 7.5, 500 mM NaCl, 0.05% Igepal, 0.5 mM EDTA) in rotation for 10 minutes at 4°C. Beads were resuspended in 40 µL of 1x Laemmli Sample Buffer (BioRad) supplemented with 50 mM DTT, and heated to 85 °C for 10 minutes; the supernatant was collected and used for Western Blot analysis, or further processed for mass spectrometry (see the “SG proteome analysis” section).

### SG RNA-Seq and bioinformatic analysis

Purified SGcore RNAs and relative inputs were retrieved in absence of doxy-induction (4 replicates); in condition of doxycycline-induced overexpression of FUS^P525L^ (2 replicates) and FUS^WT^ (2 replicates). RNA libraries for all samples were produced using Stranded Total RNA Prep with Ribo-Zero Plus (Illumina). All samples were sequenced on an Illumina Novaseq 6000 Sequencing system. Trimmomatic^71^ and Cutadapt^72^ were used to remove adapter sequences and poor quality bases; minimum read length after trimming was set to 35. Reads aligning to rRNAs were filtered out; this first alignment was performed using Bowtie2 software (v2.4.2) (https://bowtie-bio.sourceforge.net/bowtie2/index.shtml). STAR software^73^ was used to align reads to GRCh38 genome. PCR duplicates were removed from all samples using MarkDuplicates command from Picard suite (https://broadinstitute.github.io/picard/). Uniquely mapping fragments were counted for each annotated gene (ensemble release 99) using htseq-counts software^74^. edgeR software^75^ was used to compare SG enriched RNAs to relative input samples. RNAs with log_2_FC > 1 and FDR < 0.05 were defined “enriched” in SG core, while those with log_2_FC < -1 and FDR < 0.05 were defined “depleted”; all the others were labelled as “invariant”. SG from SMMC-7721^32^ cell line were re-analyzed using the previously described pipeline. For each expressed gene, the longest isoform was selected as reference. Only protein coding isoforms were taken in consideration for protein coding genes.

In the RNA-Seq analysis, the high variability of the library size of IP samples could affect the number of SG enriched RNAs identified. This could introduce technical bias especially when we need to compare the enriched RNAs of Ips performed in different conditions. To overcome this bias, we performed a “Resampling analysis” (Supplementary Fig. 2I). We decided to use the observed number of fragments in FUS^P525L^ condition experiment and then we randomly sampled fragments from the alignment files of the other conditions in order to mirror the library size composition of the reference condition. In this way, we obtained samples with similar technical variability and with a simulated similar library size. Then we compared the enriched RNAs resulted by this analysis without the bias introduced by the variability of the resulted library size. Random sampling of fragments was performed with Picard suite using FilterSamReads function and using random subsets of fragments ids list as input (https://broadinstitute.github.io/picard).

To define the SG enriched RNAs, we used the already specified log_2_FC threshold for each condition. However, the high variability of the immunoprecipitation-based experiments could result in high variability in the SG enrichment efficiency. Experiments with low enrichment efficiency could be penalized by a fixed threshold compared to others with a high efficiency, affecting the comparison of the defined enriched RNA sets. To overcome this technical limit, we performed an “Enrichment convergence analysis” (Supplementary Figure 2J). For each analyzed condition, we first ranked the expressed RNAs by their SG enrichment (log_2_FC) and then we selected a fixed number of enriched RNAs equal for each dataset. We then compared these three equally sized RNA sets using Venn Diagrams. We repeated this analysis selecting different fixed numbers of enriched RNAs, gradually reducing the inclusion of the noise from the background transcriptome (fixed numbers of top enriched RNAs: 5000, 4500, 4000, 3500, 3000, 2500, 2000, 1500, 1000, 500).

### GO term enrichment analysis

Gene ontology was performed using WebGestalt R tool (v0.4.4)^76^ applying weighted set cover reduction.

### K-mers enrichment analysis

For each analyzed condition (FUS^P525L^, FUS^WT^ and no DOXY) the SG enriched and depleted RNAs were selected. For each of the two groups the frequency of every possible k-mer of k=7 was calculated and used to compute 7-mer Z-scores. Then the difference between 7-mer Z-scores in the SG enriched group and SG depleted group was calculated and the top 50 k-mers from both sides of differences distributions were selected as top enriched or depleted k-mers in SG. For each condition, sequence logo of the top enriched 50 k-mers were computed using Logomaker Python package ^77^. After that, the average transcript frequency of each k-mer was calculated for SG enriched RNAs and background distribution (invariant group), and their ratio (fold-change, FC) was computed. Then, the log_2_FC of the top enriched and depleted k-mers of all the analyzed conditions were used for Heatmap representation and clustering performed using ComplexHeatmap R package (https://bioconductor.org/packages/release/bioc/html/ComplexHeatmap.html).

### RNA structure prediction and analysis

In RNA structure propensity analysis three indipendents random sampling of 200 mRNAs from GAIN and LOSS groups was performed imposing the similarity of 5’UTR, CDS and 3’UTR regions between groups (Mann Whitney-U Test, PValue > 0.05). RNA secondary structures were predicted using RNAfold algorithm^40^ using standard parameters and CROSS algorithm (http://service.tartaglialab.com/update_submission/736048/9393232b64) using human PARS data settings. For RNAfold prediction, the structuration score was calculated by the number of structured nucleotides over the length of the region/transcript. For RNA accessibility analysis the previously described procedure of sampling was applied to GAIN and LOSS groups, but selecting 50 mRNAs that were also detected in HEK293T cell line in the DMS-Map-Seq experiment ^42^. DMS-Map-Seq data were retrieved from RASP database^78^.

### SG Proteome analysis

After immunoprecipitation and washes (see “SG purification” section), beads were washed in 1 mL of Digestion Buffer (50 mM Sodium Bicarbonate) and resuspended in 120 µL of Digestion Buffer. Disulfide bonds were reduced on-beads by adding DTT to a final concentration of 10 mM and incubating at 56°C for 30 minutes. Proteins were alkylated by adding Iodoacetamide to a final concentration of 20 mM and incubating beads at room temperature for 20 minutes. On-beads trypsin digestion was then performed by resuspending the beads in 200 µL of Digestion Buffer supplemented with 2 µL of 0.5 µg/µL Sequencing Grade Trypsin (Merck). Samples were incubated overnight at 37°C with constant agitation at 750 rpm, then each sample (∼8 µg of peptides) was dried using a Speed-Vac and resuspended in 16 µL acetonitrile, 3% added with 0.1% formic acid. Tryptic peptides were desalted on a trapping column then loaded on a Aurora C18 column (75 mm x 250 mm, 1.6 μm particle size - Ion Opticks, Fitzroy, Australia) and separated using a Dionex Ultimate 3000 nano-LC system (Thermo Fisher Scientific). Eluents were A: water + 0.1% formic acid and B: water:acetronitrile=20:80 (*v/v*) + 0.1% formic acid. Injection volume was 2 µL, flow was 0.300 µL/min, the column temperature was 40°C, samples were eluted with a gradient program: 0.0 -1.0 min 3 % B; 1.0 – 5.0 min 3 to 5 % B; 5.0 – 35.0 min 5 to 31 % B; 35.0 – 40.0 min 31 to 44% B, 40.0 – 41.0 min 44.0 – 95 % B, 41.0 – 46.0 min 95% B and 46.0 – 47.0 min back to 3 % B. The column was then reconditioned for 13 min. The total run time was 60.0 min. Separated peptides were analyzed using an Orbitrap Exploris 480 Mass Spectrometer (Thermo Fisher Scientific), in positive ESI mode. The capillary voltage was set to 2.0 kV, the RF lens was set to 40% and the normalized AGC target % was set to 300, maximum injection time 50 ms. The acquisition was performed in Data Dependent mode (DDA) with a scan range set from 350 to 1500 m/z, resolution 120000. The Intensity threshold defined for a data dependent scan was set to 10000, MS/MS spectra were acquired in HCD mode with a collision energy of 30%. The raw data were processed using Thermo Proteome Discoverer software^79^, 1% false discovery rate (FDR) threshold and at least two peptides were used for protein identification. Proteins from Mass Spectrometry data were annotated using biomaRt (ensemble 109 release).

Protein-Protein physical Interactions were retrieved from STRING database (v11.5). Protein-RNA interactions predictions performed by catRAPID software were retrieved from RNAct database^53^. For each couple of interacting protein and RNA, the GENCODE basic isoform with the highest interaction score was selected as representative. The predicted preferential binding with GAIN over LOSS group was performed calculating for each SG protein the median normalized score with GAIN RNAs and with LOSS RNAs. Then Z-score were calculated from the ratios of median normalized scores (GAIN / LOSS) of the whole set of SG proteins.

Protein-RNA interactions analysis of experimental datasets performed using data retrieved from POSTAR3 database^54^. The preferential binding with GAIN over LOSS group was performed calculating for each RBP available in POSTAR3 database the fraction of GAIN RNAs identified as interactors of the RBP over the fraction of LOSS RNAs interactors. This analysis on GAIN and LOSS RNAs was performed considering the three matched sampling of 200 RNAs.

### Quantification and statistical analysis

The distribution and deviation of data shown in the figures of this work, the statistical tests used to calculate significant differences, and the exact value of n (e.g., number of biological replicates of the experiments) are denoted in figure legends. In main figure legends “SD” stands for “standard deviation” and “SEM” stands for “standard error mean”. Significance values were depicted in the figures using the following key legend: ∗p < 0.05, ∗∗p < 0.01, ∗∗∗p < 0.001.

