## Supplemental Figures and Legends for "ALS-associated FUS mutation reshapes the RNA and protein composition of Stress Granules"

#### Extended data Fig. 1

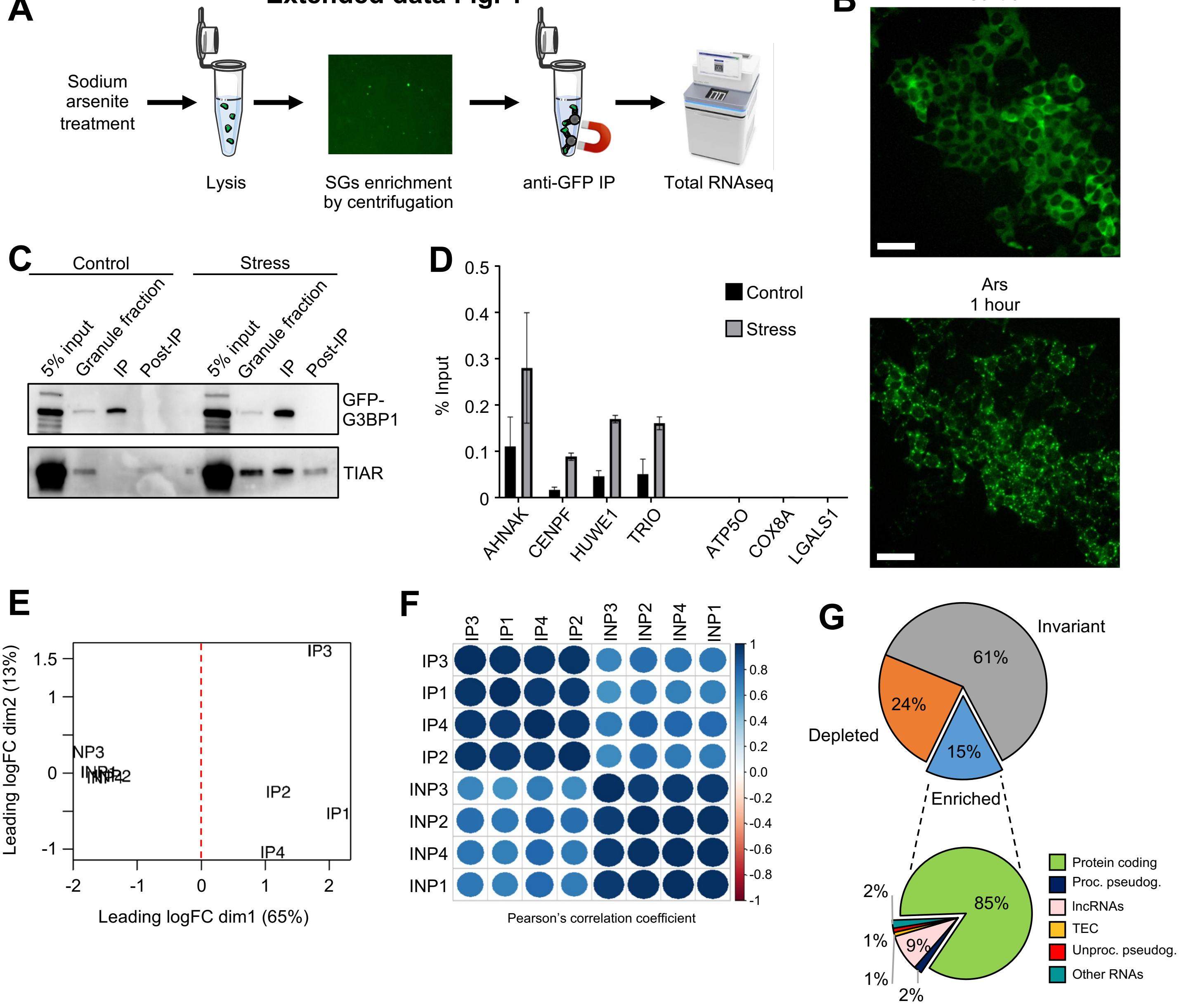


**H**

| | SK-N-BE | U2OS | SMMC-7721 |
|---|---|---|---|
| **SK-N-BE** | 1.00 | **0.82** | **0.68** |
| **U2OS** | **0.82** | 1.00 | **0.66** |
| **SMMC-7721** | **0.68** | **0.66** | 1.00 |

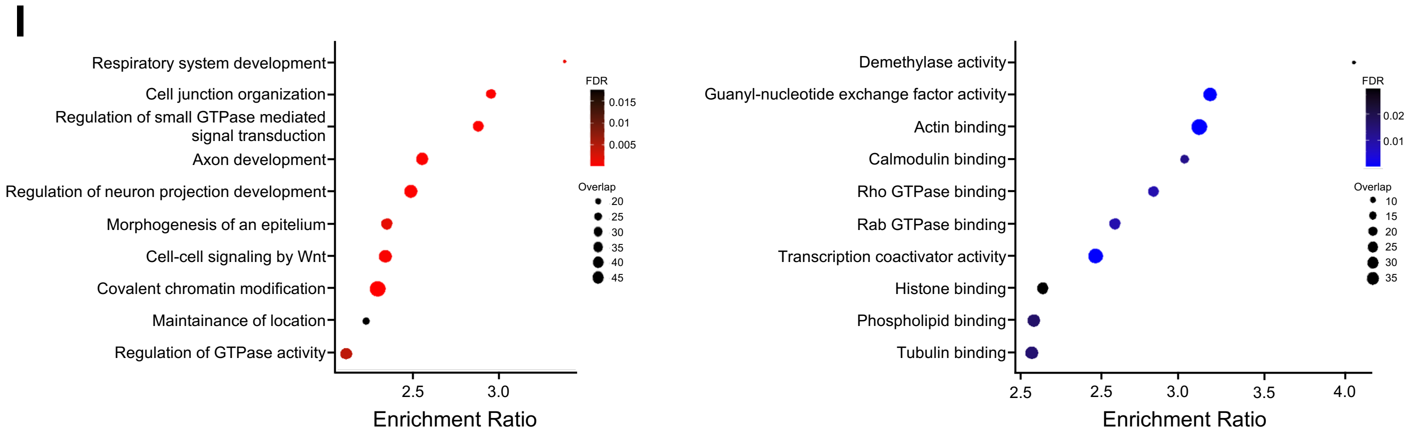

**Extended data Fig. 1**

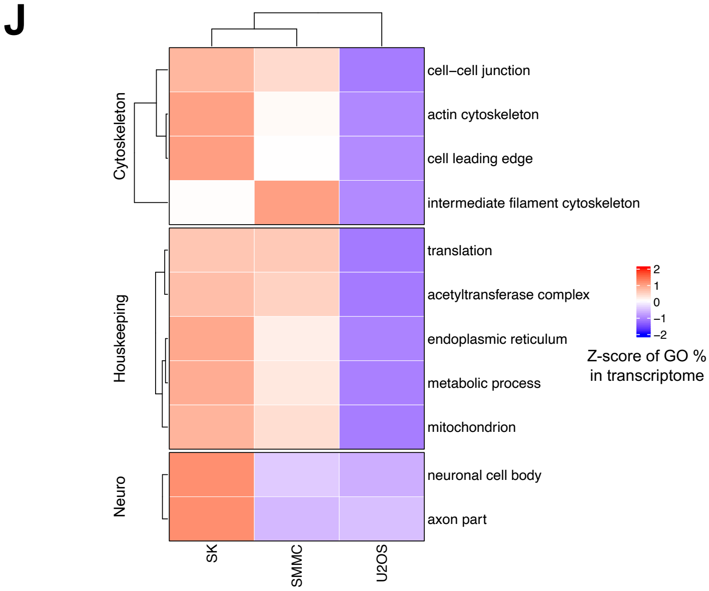


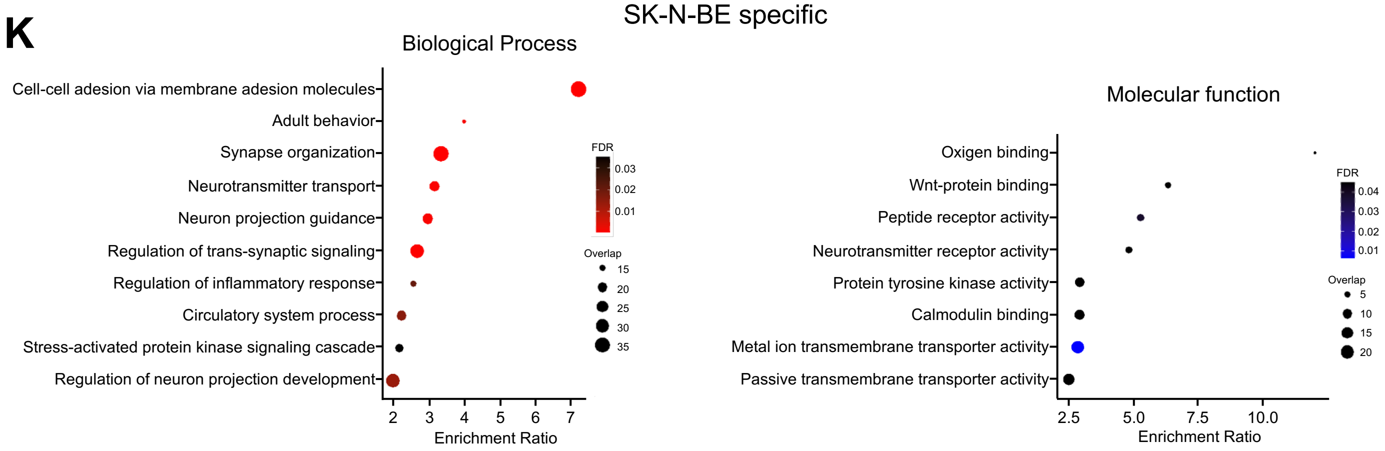


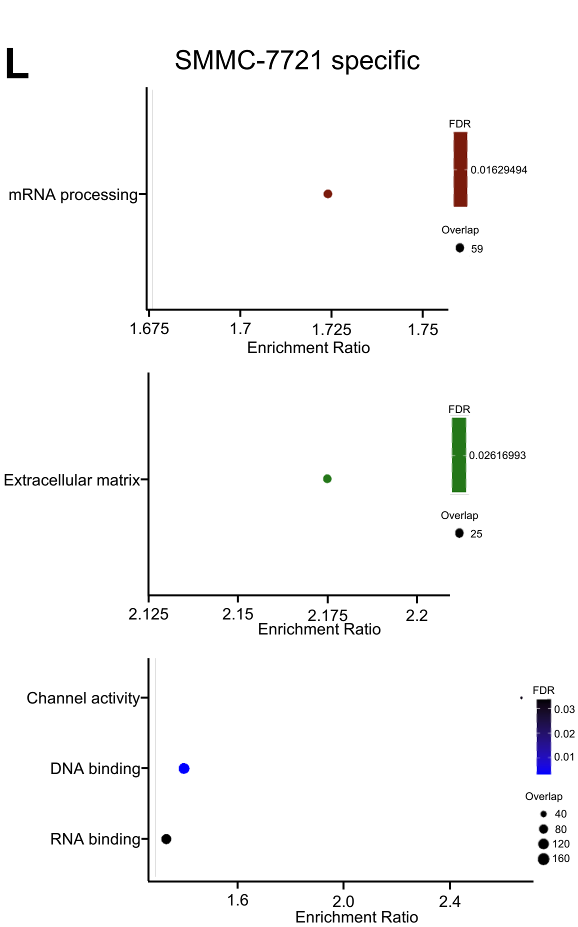


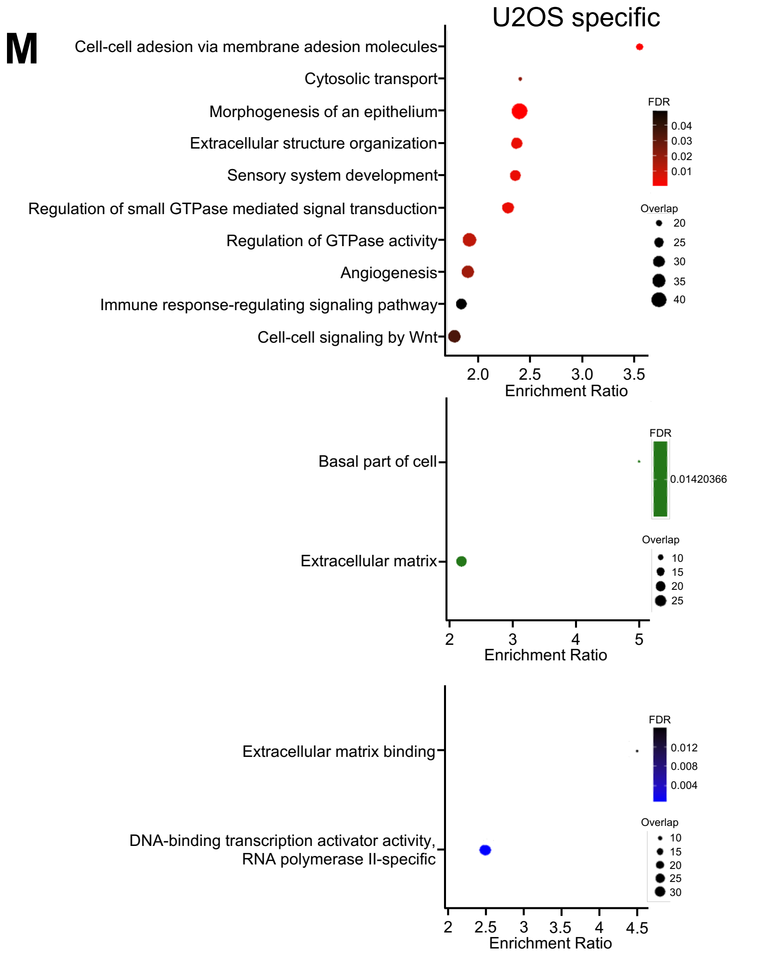

**Extended data Fig. 1**

N

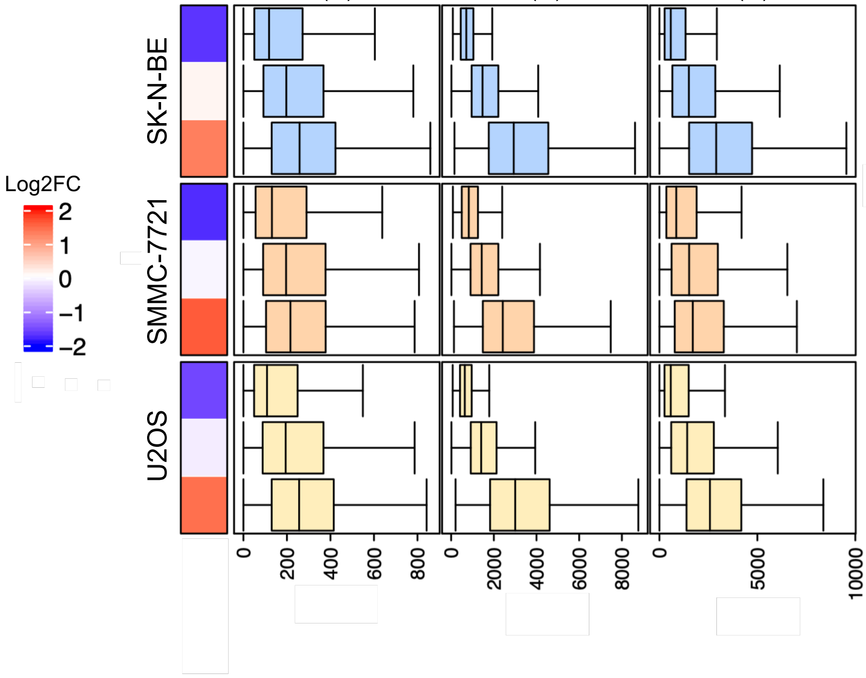


O

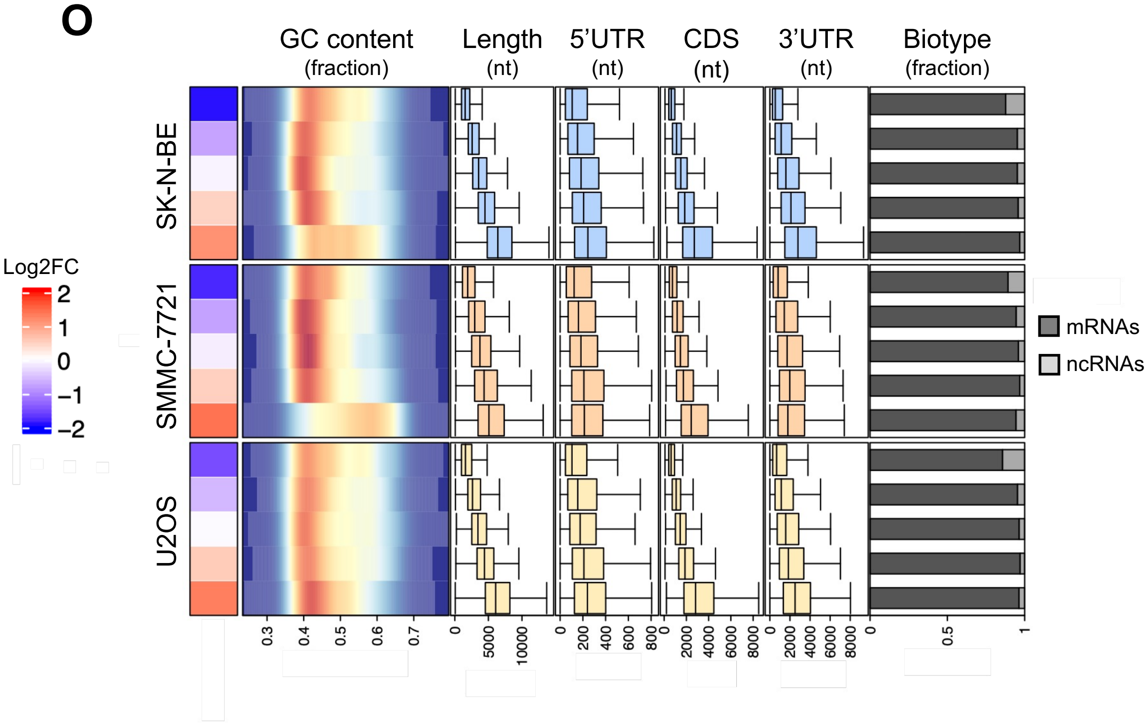

**Extended Data Figure 1**

**A)** Schematic representation of the SG purification protocol performed to isolate and analyze SG RNA composition.

**B)** Fluorescence microscopy of GFP-G3BP1 signal in SK-N-BE cells before and after oxidative stress induction with 500 mM of sodium arsenite. Scalebar = 50 μm.

**C)** Representative Western Blot analysis of GFP-G3BP1 and TIAR in SG purification samples. The Immunoprecipitation sample (IP) is compared to 5% total protein Input, the post-centrifugation fraction of the lysate ("Granule fraction"), and in the post-immunoprecipitation lysate ("Post-IP"). Control and Stress conditions were compared. TIAR was used as positive control of SG purification. n=3.

**D)** qRT-PCR analysis showing the detection of selected SG enriched (positive controls, left) and not enriched (negative controls, right) RNAs recovered from SG purification in control (black bars) or stressed SK-N-BE cells (grey bars). RNA levels are expressed as % of Input. n=3.

**E)** MultiDimentional Scaling (MDS) plot of the RNAseq samples. Each label represents a sample and the distance between 2 points reflects the leading fold change (average largest logFC between each pair of samples) of the corresponding RNAseq samples.

**F)** Correlation matrix depicting Pearson' correlation coefficients between each pairwise comparison among RNAseq samples. Positive correlations are depicted in blue while negative ones in red.

**G) Upper panel**. Pie chart depicting the proportions of SG enriched, depleted and invariant RNAs. **Lower panel.** Pie chart depicting the proportions of RNA biotypes among the SG enriched RNAs.

**H)** Table depicting Pearson's correlation coefficients among SG enrichment (log2FC) in SK-N-BE, U2OS and SMMC-7721 cell lines. Only RNAs detected in all the systems were taken in consideration for the analysis.

**I)** Dotplot depicting GO over-represented categories in the SG common *core*. Only significant categories (FDR < 0.05) of Biological Process (BP, left panel) and Molecular Function (MF, right panel) databases were depicted. X-axis represents category enrichment score while y-axis reports the GO category description. Dot size represents the amount of RNAs in the analyzed group that overlap the category while red/blue colors report the significativity of the enrichment for BP and MF categories, respectively.

**J)** Heatmap depicting the comparison (Z-score) of GO categories representation in SK-N-BE, U2OS and SMMC-7721 transcriptomes. GO categories fractions were defined as the % of GO categories genes expressed in the analyzed system. Only GO categories enriched in SG common core were analyzed. According to their biological definitions, GO categories were grouped in three macro-areas: Neuro, Housekeeping and Cytoskeletal.

**K)** Dotplot depicting GO over-represented categories in the SK-N-BE SG specific RNAs . Data are presented as in panel I.

**L and M)** Dotplot depicting GO over-represented categories in the SMMC-7721 SG specific RNAs (L) and U2OS SG specific RNAs (M). Only significant categories (FDR < 0.05) of Biological Process (BP, upper panel) Cellular Components (CC, middle panel) and Molecular Function (MF, lower panel) databases were depicted. X-axis represents category enrichment score while y-axis reports the GO category description. Dot size represents the amount of RNAs in the analyzed group that overlap the category while red, green or blue colors reports the significativity of the enrichment for BP, CC and MF categories, respectively.

**N)** Heatmap depicting the association of RNA features with SG enrichment in SK-N-BE, SMMC-7721, U2OS cell lines. For the defined transcripts group («depleted», «invariant» and «enriched») of each cell line the following characteristics were described: median log2FC (heatmap); 5' UTR, CDS and 3'UTR length (nt, boxplot).

**O)** Heatmap depicting the relationship of RNA features with SG enrichment. The 5 transcripts groups represented were defined by the stratification of the transcriptome based on SG enrichment (log2FC). For each defined transcripts group the following characteristics were represented: median log2FC (heatmap); GC content (density plot); RNA length (nt, boxplot); 5' UTR, CDS and 3'UTR length (nt, boxplot); transcripts biotypes percentages (barplot). Independent transcriptome stratifications were performed according to SG enrichment in each analyzed condition: SK-N-BE, SMMC-7721 and U2OS.

**Extended data Fig. 2**

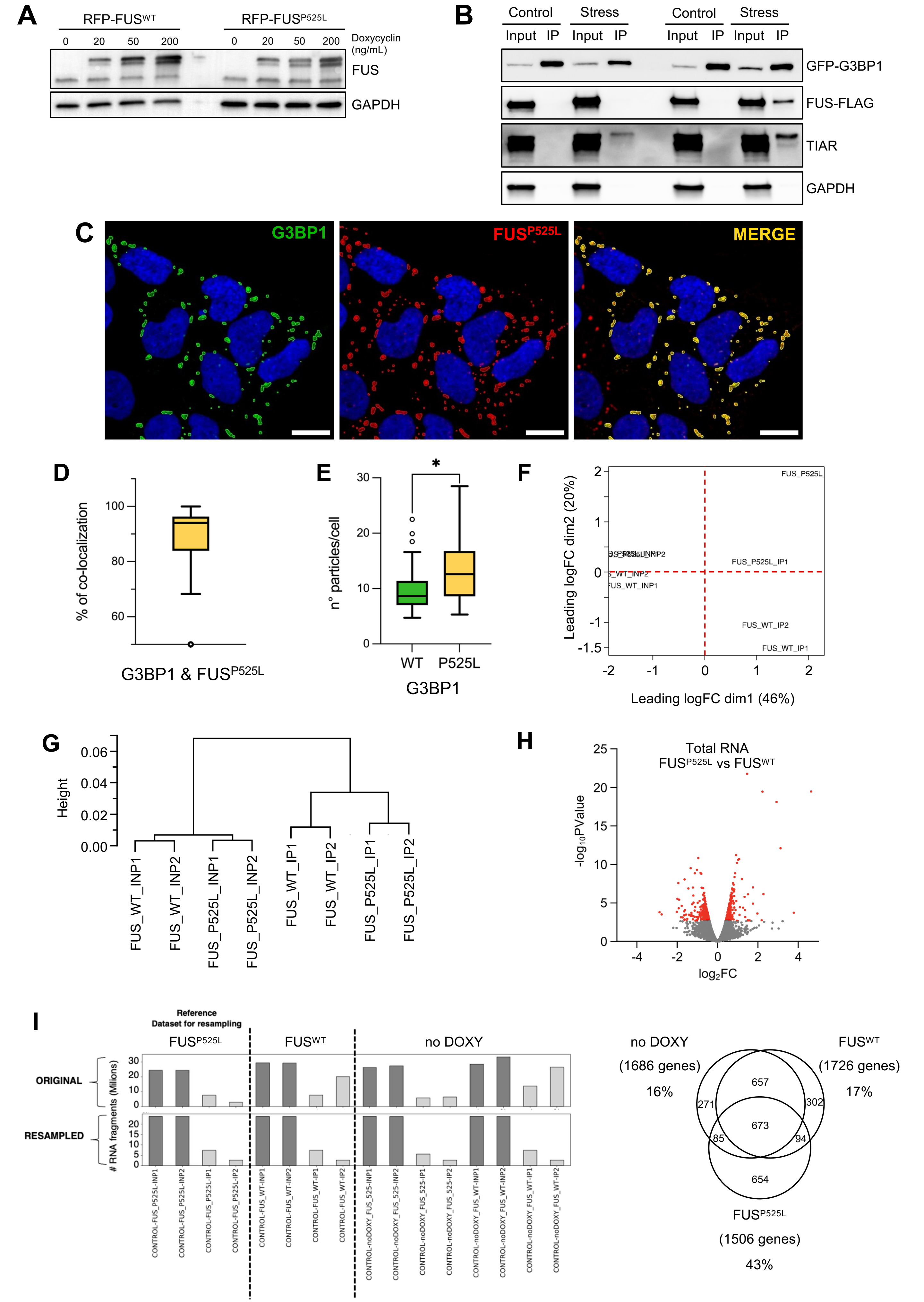

### Extended data Fig. 2

**J**

TOP 1:5000

TOP 1:500

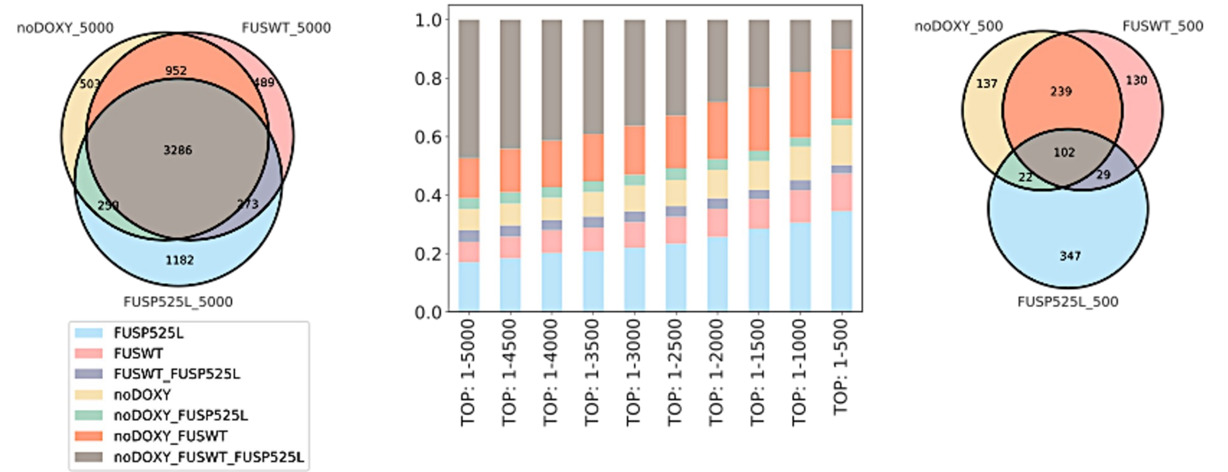


**K**

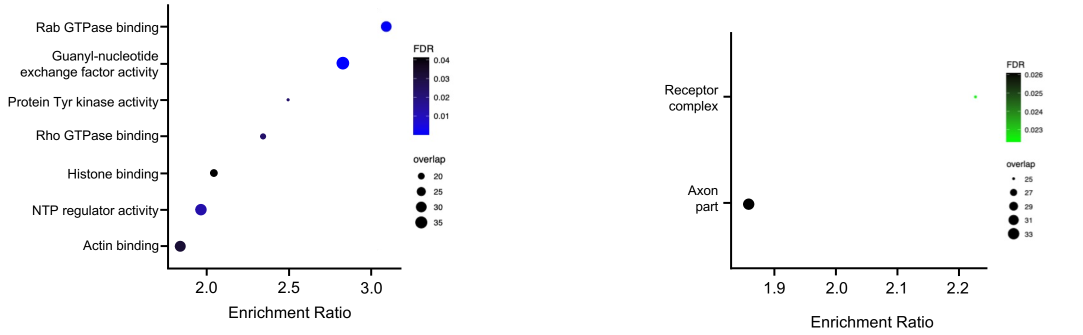


**L**

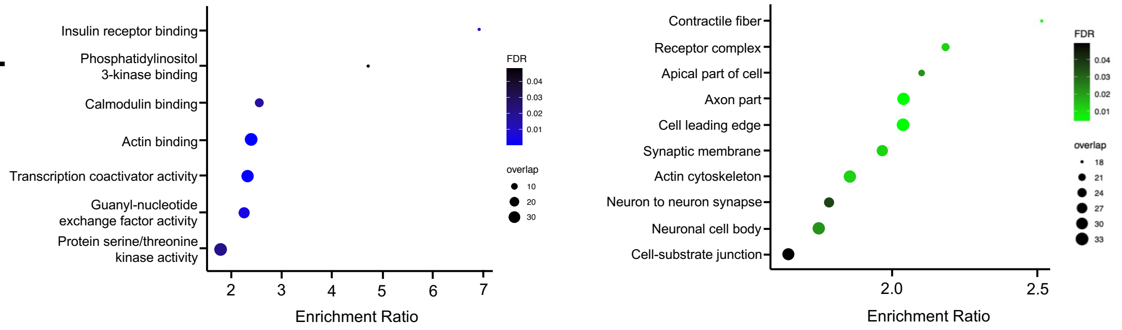


**M**

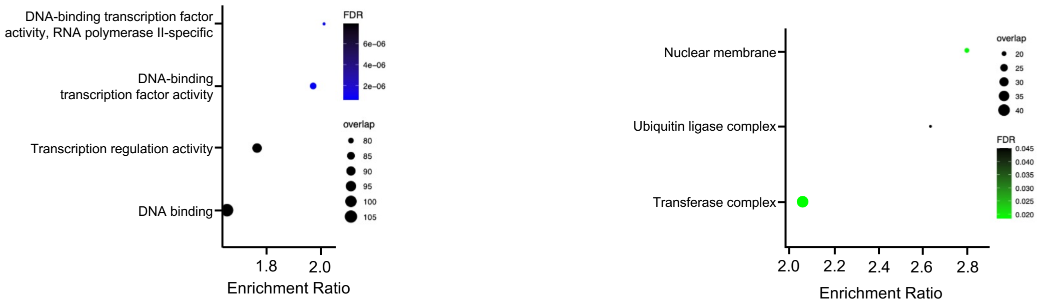

**Extended Data Figure 2**

**A)** Representative Western Blot analysis of doxycycline induction titration on SK-N-BE cells expressing an RFP-tagged form of $FUS^{WT}$ and $FUS^{P525L}$. GAPDH was used as loading control. n=3.

**B)** Representative Western Blot analysis of GFP-G3BP1 AND $FUS^{P525L}$-FLAG in samples of SG purification. The immunoprecipitation (IP) sample was compared with 5% of total protein Input (Input) and 5% of post-IP lysate (Post binding). Control and Stress conditions were compared. n=3.

**C)** Representative confocal fluorescence microscopy images of SK-N-BE cells expressing GFP-G3BP1 and an RFP-tagged form of $FUS^{P525L}$ under doxycycline control. Nuclei were counterstained with DAPI. Scalebar = 10 µm.

**D)** Boxplot indicating the percentage on colocalization of GFP-G3BP1 and RFP-FUS in SK-N-BE cells expressing the $FUS^{P525L}$ mutant protein.

**E)** Boxplot indicating the comparison of the number of GFP-positive G3BP1 particles in SK-N-BE cells overexpressing $FUS^{WT}$ and $FUS^{P525L}$.

**F)** MultiDimentional Scaling (MDS) plot of the RNAseq samples. Each label represents a sample and the distance between 2 points reflects the leading fold change (average largest logFC between each pair of samples) of the corresponding RNAseq samples.

**G)** Cluster dendrogram of Pearson's correlation distances between samples.

**H)** Volcano plot representing the differential expression analysis for RNAs detected in the Input total RNA-seq of $FUS^{P525L}$ vs $FUS^{WT}$, considering the log2 fold change ($FUS^{P525L}$ vs $FUS^{WT}$) and the -log10 p-value of expression levels. Significantly enriched or depleted RNAs (FDR < 0.05, |logFC| > 1) or invariant RNAs are indicated by red or gray dots, respectively.

**I)** **Left panel.** Barplot depicting the amount of RNA fragments in each library size of RNA-Seq samples analyzed before (upper bars) and after (lower bars) the resampling procedure. SG-enriched (light gray) and Input (dark gray) samples of $FUS^{P525L}$, $FUS^{WT}$ and no DOXY conditions were indicated. For the resampling procedure, the $FUS^{P525L}$ condition was taken as reference. **Right panel.** Venn diagrams showing the overlap between SG enriched RNAs in no DOXY, $FUS^{WT}$ and $FUS^{P525L}$ conditions after the resampling procedure. The amount of RNAs resulted enriched in SG of each condition was indicated.

**J)** Venn diagrams depict the overlap among the top 5000 (left panel) and top 500 RNAs (right panel) ranked by SG enrichment in each analyzed condition (no DOXY, $FUS^{WT}$ and $FUS^{P525L}$). Bar plot (middle panel) displays the strong divergence of $FUS^{P525L}$ SG enriched RNAs compared to the other conditions (no DOXY, $FUS^{WT}$) in every selected group.

**K, L, M)** Dotplot depicting GO over-represented categories in the $FUS^{P525L}$ COMMON (K), LOSS (L) and GAIN (M) groups. Only significant categories (FDR < 0.05) of Molecular Function (MF, blue) and Cellular Components (CC, green) databases were depicted. X-axis represents category enrichment score while y-axis reports the GO category description. Dot size represents the amount of RNAs in the analyzed group that overlap the category while green and blue color code reports the significativity of the enrichment of MF and CC categories, respectively.

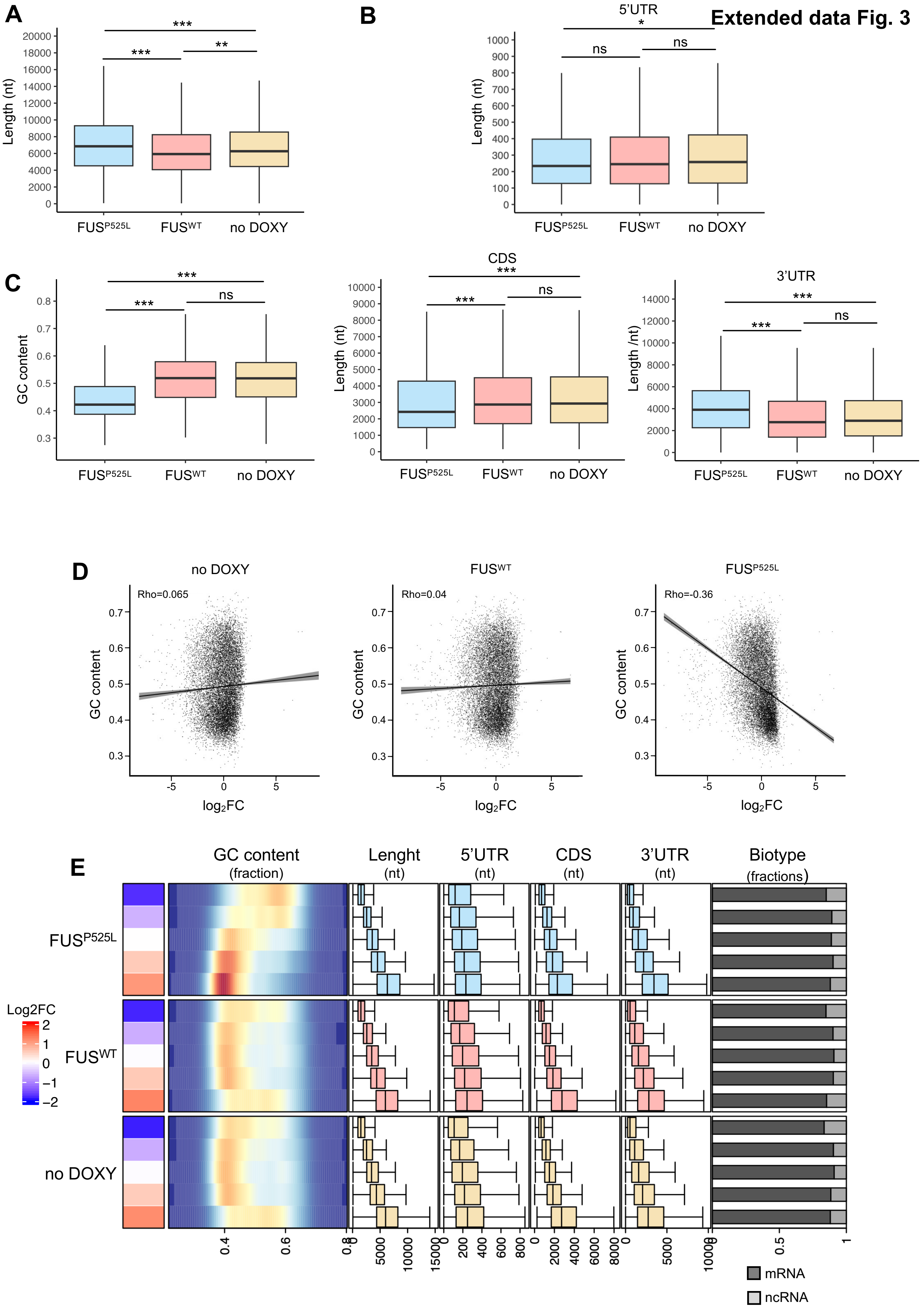
Extended data Fig. 3
A
Length (nt)
FUSP525L
FUSWT
no DOXY
B
5'UTR
Length (nt)
C
GC content
CDS
3'UTR
Length /nt)
D
no DOXY
FUSWT
FUSP525L
Rho=0.065
Rho=0.04
Rho=-0.36
GC content
log2FC
E
GC content
(fraction)
Lenght
(nt)
5'UTR
(nt)
CDS
(nt)
3'UTR
(nt)
Biotype
(fractions)
Log2FC
mRNA
ncRNA

### Extended data Fig. 3

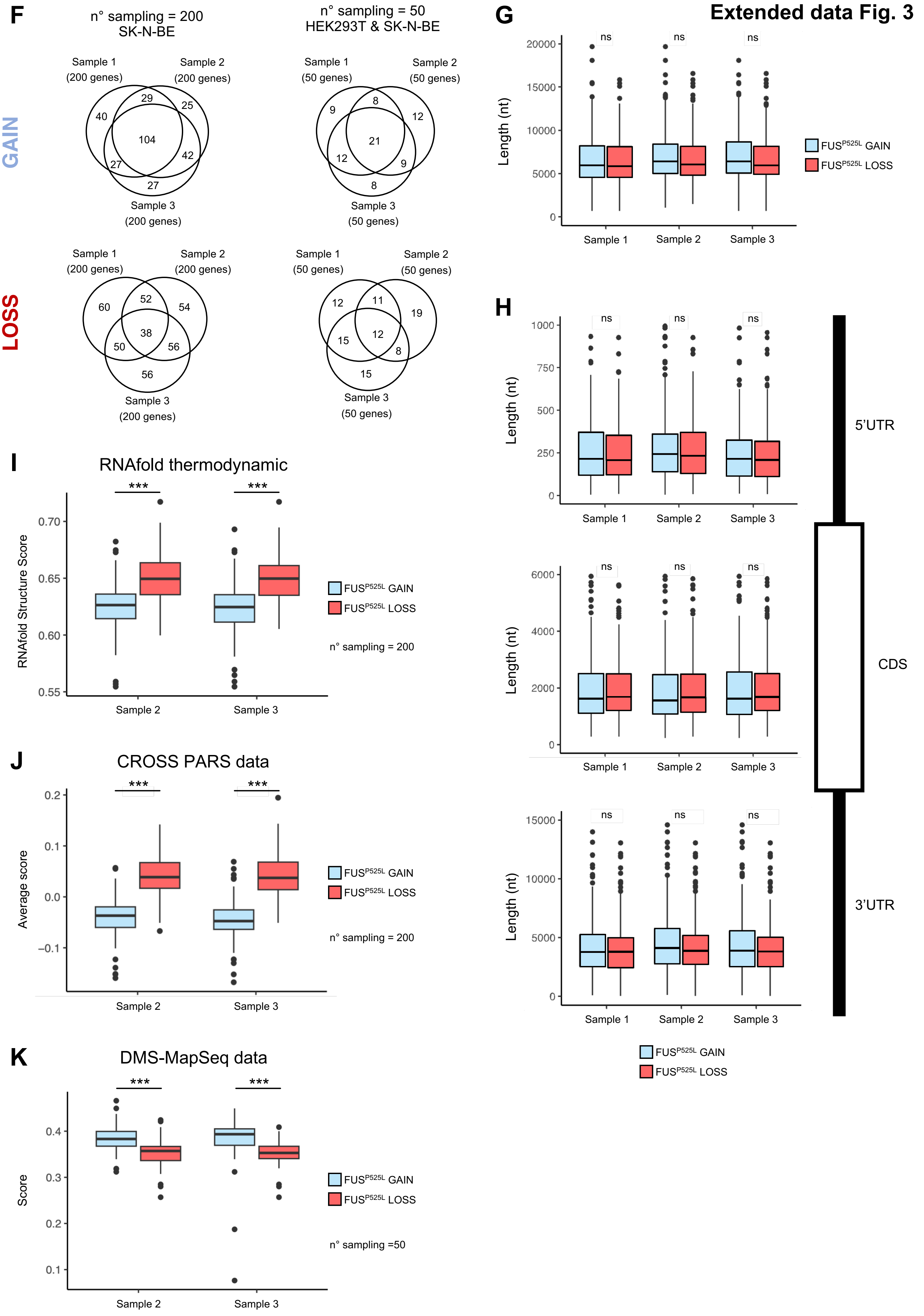

**Extended Data Figure 3**

**A, B and C)** Boxplot displaying the comparison of RNA features of the SG enriched transcripts in $FUS^{P525L}$, $FUS^{WT}$ and no DOXY condition. The following features were depicted: RNA length (A), RNA mRNA regions length (B, 5' UTR, CDS and 3'UTR) and GC content (C). Statistical significance was assessed using Wilcoxon RankSum Test.

**D)** Scatter plot showing for each RNA detected the correlation between SG enrichment (log2FC SG-enr/INP) and GC content in no DOXY (left panel), $FUS^{WT}$ (middle) and $FUS^{P525L}$ (right) conditions. Spearman' s correlation coefficients were displayed in panels.

**E)** Heatmap depicting the association of RNA features with SG enrichment. The 5 transcripts groups represented were defined by the stratification of the transcriptome based on SG enrichment (log2FC). For each defined transcripts group the following characteristics were represented: median log2FC (heatmap); GC content (density plot); RNA length (nt, boxplot); 5' UTR, CDS and 3'UTR length (nt, boxplot); transcripts biotypes fractions (barplot). Independent transcriptome stratifications were performed according to SG enrichment in each analyzed condition: $FUS^{P525L}$, $FUS^{WT}$ and no DOXY.

**F)** Venn diagrams showing the overlap between the three independent sampling of RNAs in GAIN (upper panels) and LOSS (lower panels) groups.
In the first set of sampling, 200 random RNAs with matched features (length and mRNAs regions length) were sampled from GAIN and LOSS groups (left panels).
In the second set used for DMS-Map-Seq data comparison, 50 random RNAs with matched features (length and mRNAs regions length) were sampled from GAIN and LOSS groups from RNAs expressed in both SK-N-BE and HEK293T cells and that displayed proper read coverage for DMS-Map-Seq analysis (right panels).

**G, H)** Boxplots depicting the comparable distribution of RNA lengths (G) and mRNA regions (5'UTR, CDS and 3'UTR) length (H) in the first set of 200 random RNAs sampling from $FUS^{P525L}$ GAIN and LOSS groups. The significance between groups was assessed using Mann-Whitney U test. The first, second and third sampling were displayed. Size of sampling = 200 RNAs per group.

**I, J and K)** Boxplots depicting the distribution of RNA structuration score predicted using RNAfold (I) or CROSS algorithm (J) and the distribution of RNA average accessibility calculated using DMS-Map-Seq data (K) between $FUS^{P525L}$ GAIN and LOSS groups. The differences between groups were calculated using Mann-Whitney U test. The second and third sampling were reported. Size of sampling = 200 RNAs per group for RNAfold (I) and CROSS (J) and 50 RNAs per group for DMS-Map-Seq (K).

#### Extended Data FIG. 4

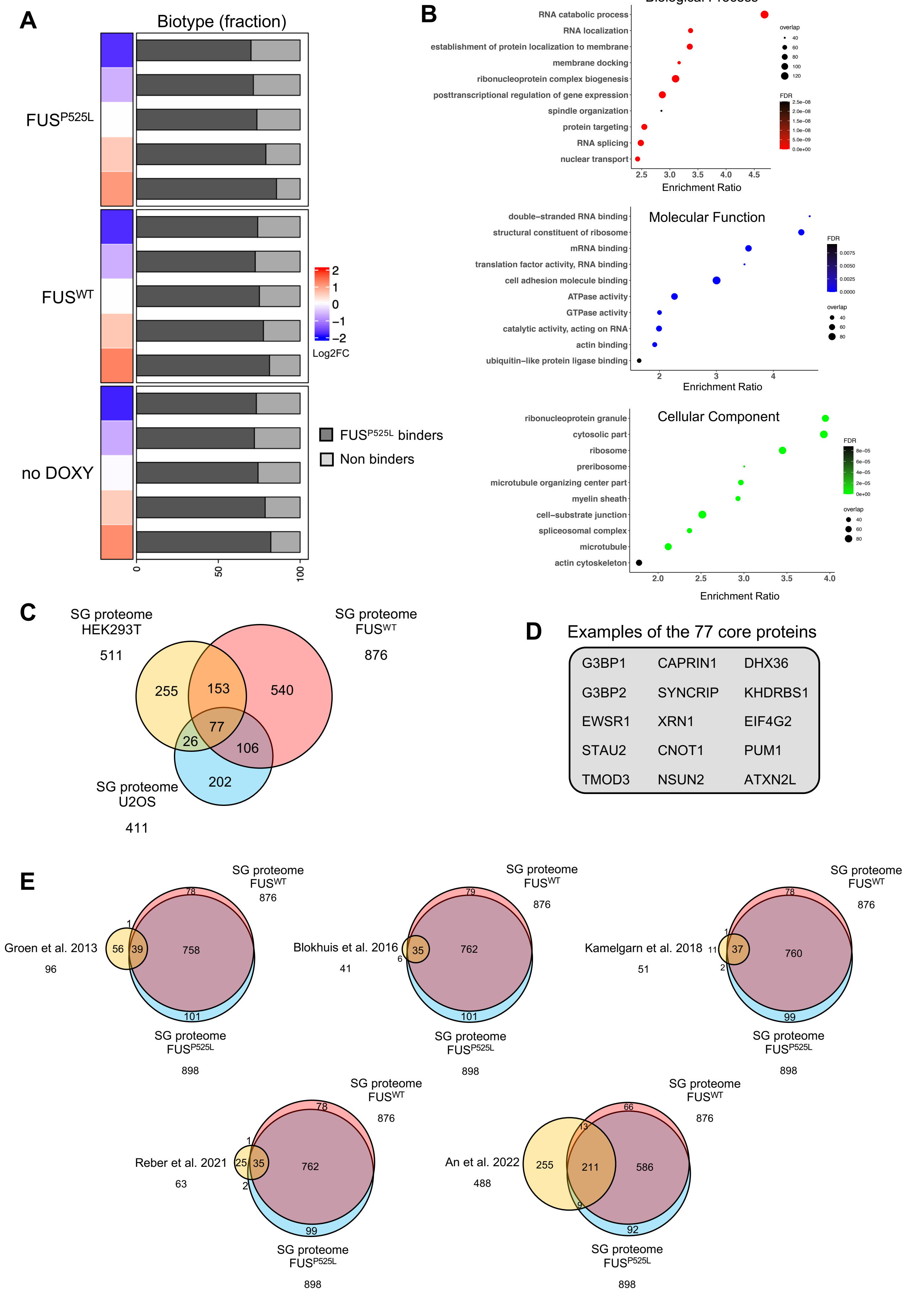

Extended Data FIG. 4

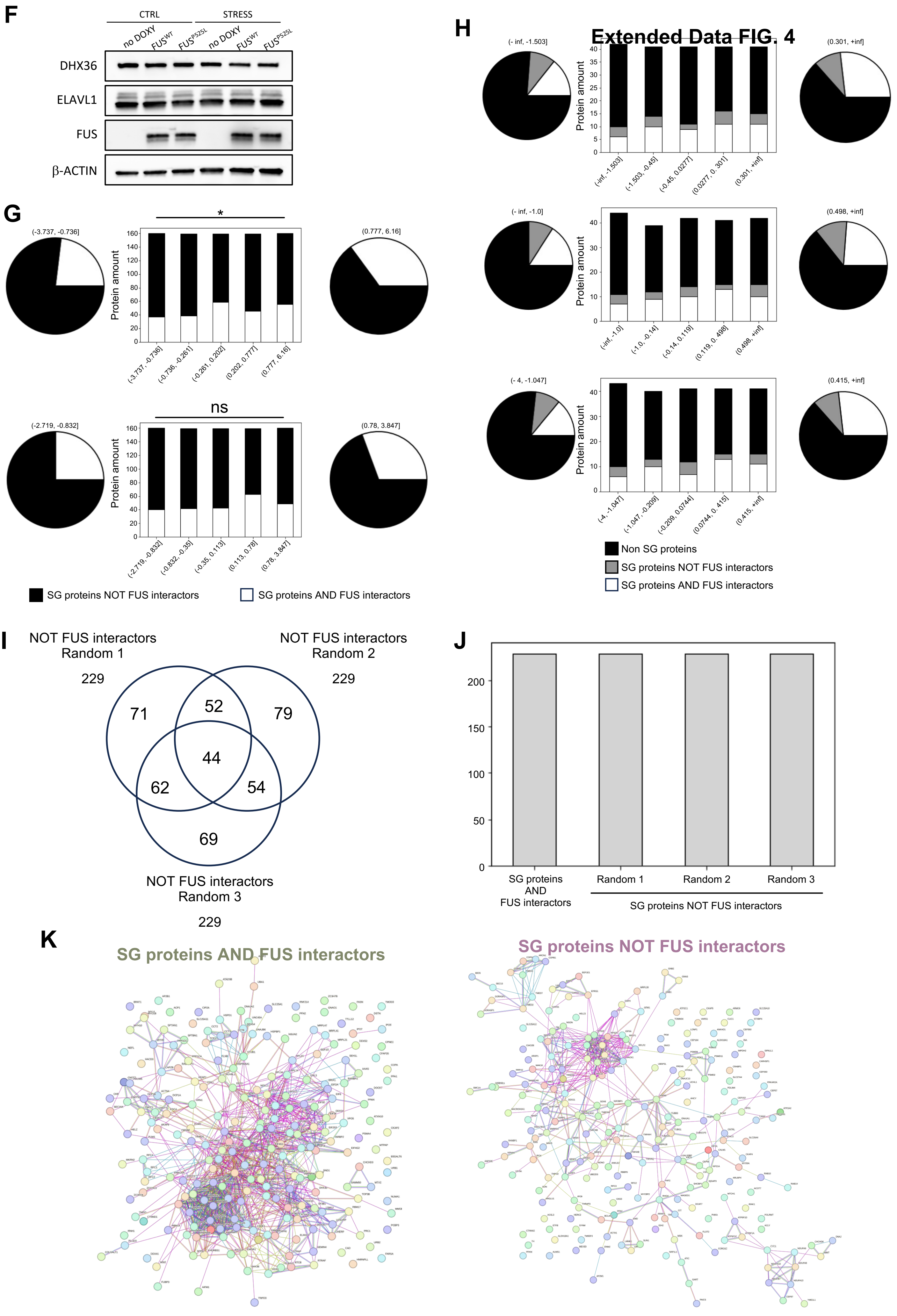

**Extended Data Figure 4**

**A)** Heatmap depicting the association between $FUS^{P525L}$ interaction with SG enrichment in $FUS^{P525L}$, $FUS^{WT}$ and no DOXY conditions. The 5 transcript groups represented were defined by the stratification of the transcriptome based on SG enrichment (log2FC). The median log2FC for each group was reported in color scale from blue to red, while barplot in the right depicts the fractions of endogenous $FUS^{P525L}$ PAR-CLIP interactors retrieved from iPSC derived motor-neurons. Only RNAs detected in both systems (SK-N-BE and MNs) were taken in consideration. Black color refers to the fraction of RNAs not bound by FUS, while white color refers to $FUS^{P525L}$ RNA interactors.

**B)** Dotplot depicting GO over-represented categories in the $FUS^{WT}$ SG proteome. Only significant categories (FDR < 0.05) of Biological Process (BP), Cellular Components (CC) and Molecular Function (MF) databases were depicted. X-axis represents category enrichment score while y-axis reports the GO category description. Dot size represents the amount of RNAs in the analyzed group that overlap the category while red, green and blue color code reports the significance of the enrichment of BP, CC and MF categories, respectively.

**C)** Venn diagram depicting the overlap of SG proteome in SK-N-BE cell in $FUS^{WT}$ condition with the proteome determined in U2OS and in HEK293T cells.

**D)** Selected examples of proteins that compose the SG common *core*.

**E)** Venn Diagrams depicting the overlap of published FUS protein interactors and the SG proteome described in this work.

**F)** Representative Western Blot analysis of DHX36, ELAVL1 and FLAG-FUS in the Input fractions of the SG purification experiments in Fig. 4C. β-Actin was used as loading control. n=3.

**G)** Results showing the analysis on RNA-protein interaction made with catRAPID algorithm. Pie charts depict the fraction of ”SG protein AND FUS interactors” (white) and ”SG proteins NOT FUS interactors” (black) in the last (left) and first group (right), while the Bar plot (middle) displays the indicated fraction in every stratification group. For GAIN and LOSS comparison the matched length sampling were used. The second and third sampling were reported. Significance of the enrichment of ”SG proteins AND FUS interactors” in the first group compared to the last was assessed with Fisher’s exact test.

**H)** Results representing experimental data of RBP-RNA interactions from POSTAR3 database. Pie charts depict the fraction of ”non SG proteins” (black), ”SG protein AND FUS interactors” (white) and ”SG proteins NOT FUS interactors” (grey) in the last (left) and first group (right), while the Bar plot (middle) displays the indicated fraction in every stratification group. For GAIN and LOSS comparison the matched length sampling were used. The three samplings were reported (upper, middle and lower panels respectively). Significance of the enrichment of ”SG proteins AND FUS interactors” in the first group compared to the last was assessed with Fisher’s exact test.

**I)** Venn diagram depicting the overlap between the down-sampled groups of proteins randomly selected from the ”SG protein NOT FUS interactors” group.

**J)** Barplot depicting the comparable amount of proteins in “SG protein AND FUS interactors”, and the three down-sampling groups of proteins randomly selected form the ”SG proteins NOT FUS interactors” group.

**K)** Network of protein-protein physical interactions in “SG proteins AND FUS interactors” and in one of the three sampling performed from the “SG protein NOT FUS interactors group”, retrieved from STRING database.
